# Membrane proximal F-actin restricts local membrane protrusions and directs cell migration

**DOI:** 10.1101/705509

**Authors:** Anjali Bisaria, Arnold Hayer, Damien Garbett, Daniel Cohen, Tobias Meyer

## Abstract

A major component of cell migration is F-actin polymerization driven membrane protrusion in the front. However, F-actin proximal to the plasma membrane also has a scaffolding role to support and attach the membrane. Here we developed a fluorescent reporter to monitor changes in the density of membrane proximal F-actin during membrane protrusion and cell migration. Strikingly, unlike total F-actin concentration, which is high in the front of migrating cells, the density of membrane proximal F-actin is low in the front and high in the back. Furthermore, local membrane protrusions only form following local decreases in membrane proximal F-actin density. Our study suggests that low density of membrane proximal F-actin is a fundamental structural parameter that locally directs membrane protrusions and globally stabilizes cell polarization during cell migration.

**One Sentence Summary:** Membrane protrusion and cell migration are directed by local decreases in the density of membrane proximal F-actin

## Introduction

The cell cortex is a complex actin meshwork below the plasma membrane of all eukaryotic cells (*1*). Changes in its organization, thickness, and contractility determine cell shape dynamics during mitosis, blebbing and other processes (*2*). The actin cortex also acts as a scaffold that allows transmembrane “pickets” to organize and cluster transmembrane and surface receptors by altering their diffusion-reaction dynamics (*3*, *4*, *5*). This coupling may be altered by the distance and/or density of the most proximal cortical F-actin network. Furthermore, the F-actin density below the membrane regulates membrane adhesion, and consequently membrane tension, by providing more substrate for ezrin, radixin and moesin family (ERM) and other membrane tethering proteins (*6*). Despite the fundamental relevance of the actin cortex, our understanding of membrane proximal F-actin (MPA) density as a local structural parameter involved in cell migration and other cellular processes remains limited.

Recently STED microscopy in living cells found spatial variations in the actin cortex, suggesting that it is locally regulated (*7*). This can be explained by signaling mechanisms that locally regulate polymerization, severing and capping of membrane proximal F-actin as well as changes in the activity of ERM and other linker proteins that attach the actin cortex to the membrane (*8*). Tether experiments that measure membrane tension showed that global reduction in MPA density reduces membrane tension, increases bleb formation in fibroblasts (*9*) and, in migrating zebrafish mesoderm cells, impedes persistent migration (*10*). Furthermore, cycles of membrane tension may help specify the leading edge through long range mechanical signaling (*11*). *A priori* one would expect that the front of a cell should have an increased MPA density due to the enrichment of F-actin in lamellipodia of cells migrating in both 2D and 3D (*12, 13*). If true, however, this would lead to increased attachment between the membrane and F-actin which would oppose protrusion at the front. Nevertheless, previous studies also found higher ERM phosphorylation and activity at the rear of migrating amoeboid cells and leukocytes (*14*–*17*), suggesting that MPA density might also be high in the back. Thus, regulation of the local density of MPA could have a fundamental but as of yet poorly understood role in cell migration and other cellular processes that depend on cortical actin. We hypothesized that migrating cells might have spatial gradients and local differences in MPA density which could in turn alter membrane tension, membrane cytoskeleton attachment, and protrusion dynamics. To test this, an unbiased measure of the local density of MPA is needed that avoids the side effects resulting from overexpressing biologically-active scaffolding proteins like ERMs or relying on site-specific techniques such optical tweezers to pull membrane tethers.

Here we develop a fluorescent reporter that measures spatial variations in the membrane proximal F-actin density using standard microscopy techniques. Using this <u>M</u>embrane <u>P</u>roximal F-<u>Act</u>in (MPAct) reporter, we uncovered robust front-to-back gradients of increasing F-actin density in the membrane proximal attachment zone in polarized, migrating cells. Strikingly, the density of MPA is low towards the front of migrating cells despite an opposing high concentration of F-actin in the same front region. Similar gradients were observed in single and collectively migrating cells and in cells migrating in 2D and 3D environments. Surprisingly, we find that areas with high MPAct signals do not respond to increases in Rac1 activity, but protrusions are instead biased towards areas with low MPAct signals. We conclude that the density of MPA is a key structural parameter that is locally reduced to direct local protrusions, and that gradients of MPA stabilize cell polarity to promote persistent cell migration.

## Results

### A fluorescent live-cell reporter that measures the density of membrane proximal F-actin

Consistent with results in amoeboid migration (*14*–*17*), we observed an enrichment of fluorescent protein-conjugated moesin in the rear of collectively migrating hTERT immortalized human umbilical cord vein endothelial cells (HUVEC) and individual Retinal Pigmented Epithelia (RPE-1) cells (Fig. 1A-B, Movie S1). Nevertheless, siRNA knockdown of ezrin or moesin, the two main ERM proteins expressed in HUVEC, (Fig. S1A), did not show significant effects on cell speed or persistence in these cells, suggesting that multiple direct and indirect mechanisms may link the plasma membrane to the actin cortex (Fig. 1C, S1B-D). Notably these mechanisms include: local PIP_2_ levels modulate actin severing and capping, small GTPases affect Arp2/3 and formin-mediated actin polymerization, ROCK and PAK regulate contractility, and localization of ERMs and other scaffolding proteins (e.g. EBP50 or E3KARP) reinforce membrane attachment (*18*). This multiplicity of scaffolding and regulatory mechanisms of cortical actin motivated us to instead take a structural approach to design a fluorescent reporter system that measures local density of MPA independent of particular attachment proteins or regulatory mechanisms.

**Fig. 1.**
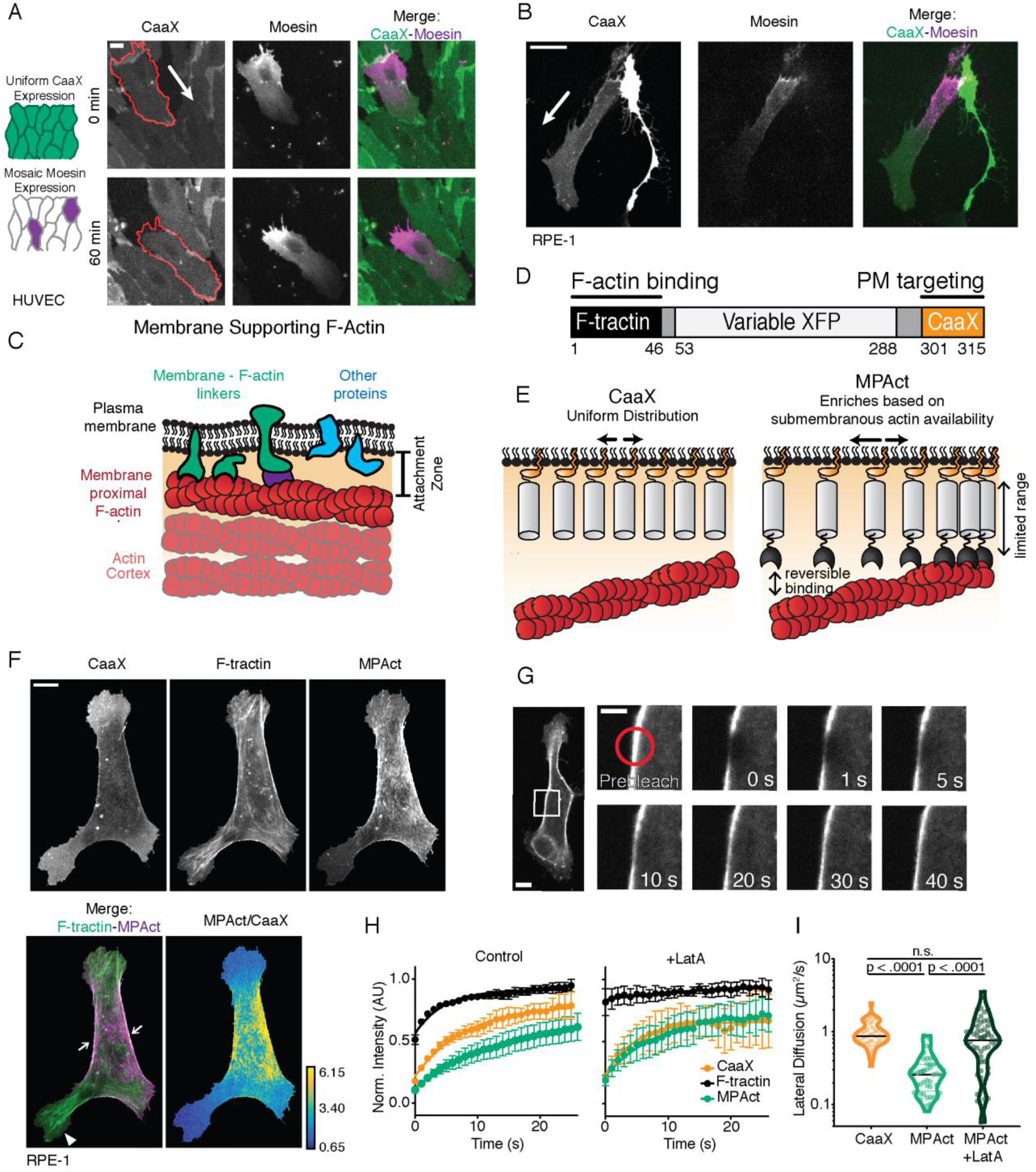
A fluorescent reporter for cortical actin accessibility reversibly binds to membrane proximal F-actin. (**A**) Monolayers expressing iRFP-CaaX (left, green) transiently transfected with GFP-moesin (middle, magenta). Cell outline in monolayer shown in red. (**B**) iRFP-CaaX (left, green) and GFP-moesin (middle, magenta) transiently co-expressed in randomly migrating RPE-1 cells. Direction of migration indicated by arrow. (**C**) F-actin in a membrane proximal “attachment zone”, part of much thicker actin cortex, acts as substrate for membrane-F-actin attachment (by ERMs and other mechanisms). (**D**) Design of membrane proximal actin density (MPAct) reporter, combining a CaaX membrane anchor from K-Ras4B and actin binding motif F-tractin connected by two short linkers and different fluorescent proteins (FP). (**E**) Schematic of expected locally increased distribution for MPAct (right) versus CAAX density in areas with higher MPA density. (**F**) RPE-1 cells transiently co-transfected with MPAct-mCitrine (magenta), CFP-CaaX, and F-tractin-mCherry (green). CaaX normalized MPAct signal as parula colormap where yellow is the highest and blue lowest ratio. (**G**) (Right) Image series showing recovery after local photobleaching of lateral membrane MPAct in RPE-1. (bar = 5 μm for right images). (**H**) FRAP analysis in HeLa of F-tractin-mCitrine (Black), YFP-CaaX (Orange), or MPAct-mCitrine (Green) under control conditions (left) or after treatment with 1 μM of Latrunculin A (LatA, right). n = 10 cells for all conditions. Error bars are 95% confidence intervals (CI). (**I**) Diffusion rates calculated from FRAP experiments in RPE-1 using 3D Gaussian profile fitting for CFP-CaaX, and MPAct-mCitrine with and without 1 μM LatA. n=52,48, and 54 respectively, 2 independent experiments. Dotted lines are quartiles. p-values from unpaired t-test. Scale bar is 10 μm unless marked.

We designed a reporter to measure the local density of F-actin specifically in the attachment zone by using a fast-diffusing plasma membrane anchor that is linked to a fluorescent protein and a low-affinity F-actin binding domain (Fig. 1D). Our rationale was that the reporter will dynamically equilibrate and relatively increase its concentration in regions of the plasma membrane with high MPA density. As a membrane anchor, we used a polybasic farnesylated CaaX motif derived from K-Ras4B (*19*), which selectively localizes to the plasma membrane and quickly diffuses at ~1 μm^2^/s along the inner membrane leaflet (*20*–*22*). We used a small reversible F-actin binding domain of inositol 1,4,5-trisphosphate 3-kinase (F-tractin) (*23*) which has been used as a live-cell marker of all F-actin in cells (*24*–*26*). F-tractin has a low ~3 μM binding affinity for F-actin, and is more reversible and less interfering compared to other commonly used F-actin markers such as utrophin or life-act (*27*).

The MPAct reporter system is based on ratiometric analysis of the relative local concentration of MPAct compared to a plasma membrane marker with the same CaaX motif. Due to its fast diffusion, the MPAct signal reports relative enrichment and depletion of MPA concentration in different regions of the cell cortex relative to the membrane marker (Fig. 1E). The direct comparison to a membrane marker is critical for the analysis due to local submicroscopic changes in membrane architecture (e.g. invaginations) that can make increased membrane concentration appear as increased MPAct signal. Short linkers ensure that the reporter monitors the MPA density in the attachment zone within less than 10 nm from the membrane surface. In addition, the low affinity and rapidly reversible binding of F-tractin ensures that fast changes in the actin cortex can be monitored as the actin cortex is continuously turned over with a half time of 20-25 seconds (*28*).

We first tested if the MPAct reporter localizes differently from its component parts – F-tractin or CaaX alone (Fig. 1F). Confocal imaging of RPE-1 cells transiently co-transfected with fluorescently labeled CaaX, MPAct, and F-tractin revealed that the MPAct has a different distribution from the CaaX membrane marker. Furthermore, while many of the regions where the reporter was enriched were also enriched in F-tractin (Fig.1F arrows), other regions had high F-tractin signals but low MPAct signals, implying that some F-actin rich regions can have a low MPA density (Fig. 1F, arrowhead). These low MPAct regions can be readily identified after normalizing with the CaaX membrane marker (Fig 1F, bottom right) as large blue areas rich in F-tractin. Similar results were seen in HeLa cells (Fig. S2A) and control experiments using phalloidin staining showed that the reporter does not cause observable changes to F-actin structures (Fig. S2B).

We next determined whether the reporter is binding membrane-proximal F-actin by measuring lateral membrane diffusion using fluorescence recovery after photobleaching (FRAP) analysis (Fig. 1G). We bleached small areas (< 4 μm^2^) of the lateral membrane in HeLa transiently expressing CaaX, F-tractin, or MPAct fused fluorescent proteins and monitored recovery over the course of 30 seconds. As expected, cytosolic F-tractin showed a rapid recovery (Fig. 1H, black line t_½_= 2.32 ± 0.38 s), the CaaX construct showed a slower recovery and MPAct the slowest (t½= 5.50 ± 0.97s (orange line) and 9.54 ± 2.70 s (green line) respectively, n > 10 for all conditions). After addition of 1 μm Latrunculin A (LatA), a potent F-actin inhibitor, the recovery kinetics of MPAct became nearly identical to the CaaX marker (t_½_= 3.31 ± 1.01s and 4.51 ± 1.60 s respectively). We also verified that this difference was due to a difference in lateral diffusion by explicitly fitting the initial recovery after FRAP to a Gaussian diffusion profile as described previously (Fig. S2C (*22*)). CaaX alone showed a lateral diffusion of 0.96 ± .05 μm^2^/s which is similar to previously reported values (Fig.1I). MPAct showed a slower lateral diffusion than CaaX under normal conditions (0.30± .03 μm^2^/s), but this difference was abolished after addition of 1 μM LatA (0.88 ± .1 s μm^2^/s). Similar lateral diffusion rates results were found in RPE-1 cells (Fig. S2D). Finally, we constructed a MPAct mutant with an F-tractin domain deficient in binding F-actin (MPAct(Δ1-6) Fig. S2E-F) and found that this construct has a fast lateral membrane diffusion similar to the CaaX construct (Fig. S2F (*23*)). Thus, the slower diffusion of MPAct relative to the CaaX probe is caused by reversible binding of the MPAct reporter to MPA.

### Stably polarized cells migrating on linear tracks have gradients with low MPAct in the front

To determine if migrating cells have a high MPA density that corresponds to high F-actin concentration in lamellipodia, we used hTERT HUVEC expressing differentially tagged CaaX, F-tractin, and MPAct (*24*). Cells were plated on linear 20 μm wide fibronectin tracks surrounded by non-adhesive poly-L-lysine polyethylene glycol (PLL-PEG) to promote persistent polarization and migration (Fig. 2A, described in (*29*)). Strikingly, the MPAct reporter revealed a gradient of MPA that is lowest in the front - even though we confirmed that the F-actin concentration is high in the front 3 μm region of these cells using F-tractin (representative cell shown in Fig. 2B). These MPAct gradients were stable over 60 minutes while cells persistently migrated (Fig. 2B-C, Movie S2) based on a kymograph analysis described Fig. S3A. We conclude that MPA is highly polarized along the axis of migration with low levels in the front in 1D migration. We found similar gradients in RPE-1 cells plated on 20 μm fibronectin tracks (Fig. S3B). Though this result was unexpected, we also found some areas enriched in MPAct colocalize with areas of high moesin (Fig. 2D), consistent with previous conclusions that ERMs mark areas of increased MPA density. Yet, siRNA knockdown of ezrin or moesin causes no change in the observed MPAct gradient, suggesting that multiple mechanisms likely contribute to establishing an MPA density gradient in migrating cells (Fig. S3N).

**Fig. 2.**
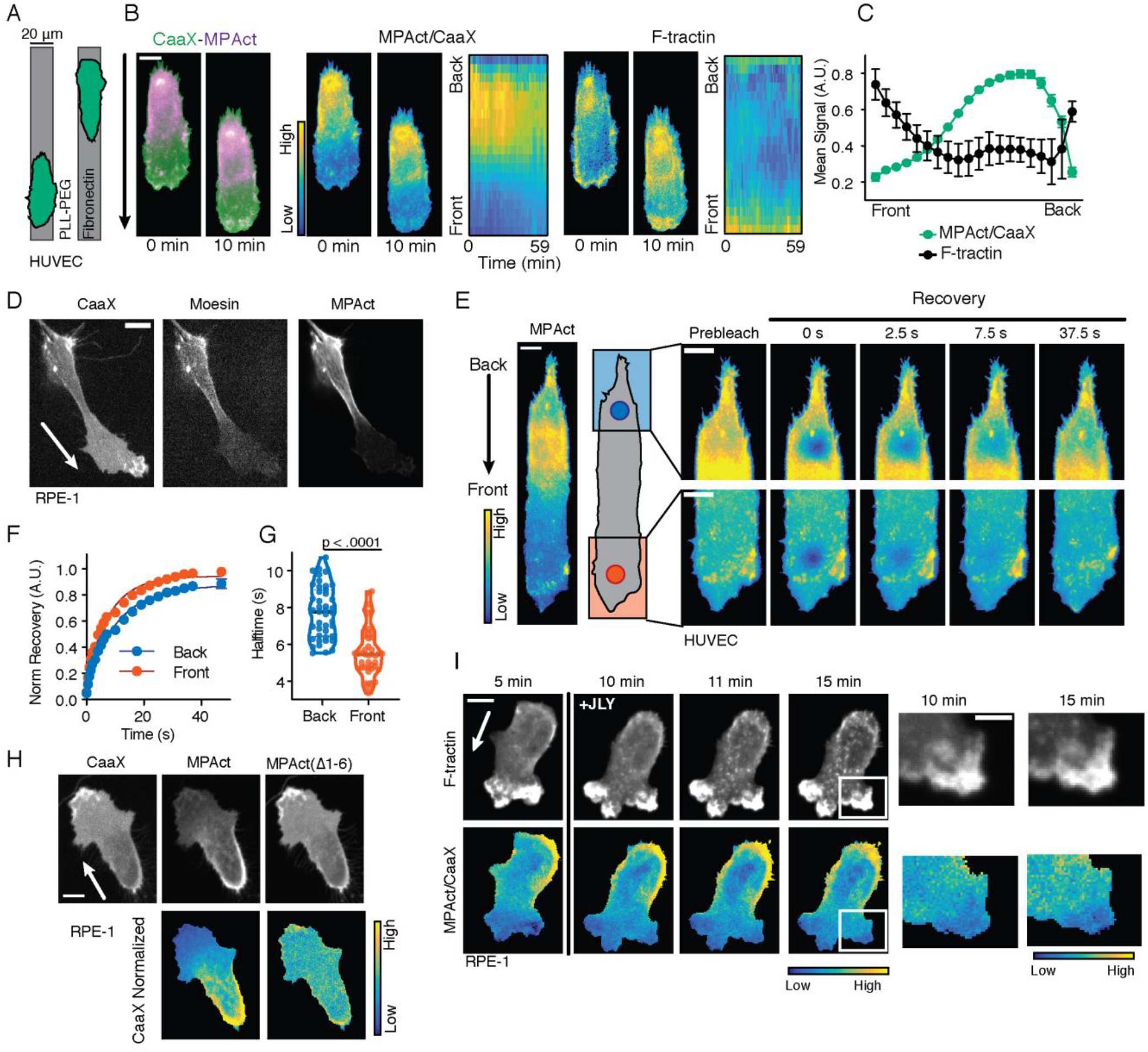
Migrating cells show stable back-to-front gradients with low membrane proximal F-actin density in the front. (**A**) Schematic of fibronectin stripes stamped using PDMS molds. PLL-PEG is used to create non-adherent regions. (**B**) (left) HUVEC stably expressing MPAct-mCitrine (magenta) and iRFP-CaaX (green) migrating on stripes. Ratio of MPAct/CaaX (middle) and F-tractin signal (right) at same time points. Kymographs of activity changes – front-to-back location versus time are also included. (**C**) Quantification of MPAct and F-tractin activity profiles over time for cell in (B) as an example. Error bars are standard deviation. (**D**) Randomly migrating RPE-1 transiently expressing iRFP-CaaX, MPAct-mRuby3, and GFP-moesin. (**E**) Photobleaching recovery of MPAct-mCitrine in HUVEC migrating on 2D fibronectin stripes comparing front versus back. (**F**) Average recovery curves, N=48 matched cells from 3 independent experiments. (**G**) Halftimes calculated from individual recovery curves in E. p-values from paired t-test. (**H**) (Top) RPE-1 cells co-expressing CFP-CaaX, MPAct-mCitrine, and MPAct(Δ1-6)-mRuby migrating randomly on collagen coated glass. (Bottom) MPAct and MPAct(Δ1-6) normalized to CAAX reference are shown below. (**I**) RPE-1 treated with actin-arresting JLY cocktail described in and Methods section. Time course of F-tractin-mCherry (top) and MPAct signals (bottom). “Frozen” protrusion highlighted in box shown expanded on the right (bar = 5 μm). For all panels direction of cell migration indicated by arrow. Scale bar is 10 μm unless marked. Scale of MPAct/CaaX varies with image modality and cell shape and is normalized to show maximal signal range.

We then used an orthogonal method to test whether MPAct polarity reports a lower level of MPA in the front. We employed fluorescence recover after photobleaching (FRAP) analysis by bleaching identical ~5 μm^2^ areas in the front and back of cells migrating on fibronectin stripes and compared the recovery kinetics (Fig. 2E, S3D). Because the reporter shows slower recovery when bound to F-actin (Fig 1I), we surmised that recovery rates in different regions would reflect the amount of F-actin MPAct had access to. Consistent with this expectation, the back of migrating cells showed slower recovery (t_½_ = front: 5.48 ± 0.39 s and back: 7.53 ± .605 s) when compared to the front of cells (Fig. 2F, G; also S3E) and also by analyzing the recovery kinetics in the front and back of randomly migrating RPE-1 (Fig. S3F). We additionally used an MPAct(Δ1-6) F-actin binding mutant to show that F-actin binding is required to create the gradient (Fig. 2H). Together, these results further support our interpretation that MPA density is reduced in the front of cells.

We next addressed a concern that actin treadmilling or other dynamic processes in the front of cells may prevent full equilibration and exclude the MPAct reporter. We considered that during a protrusion, a CaaX marked membrane might rapidly extend, while the slower diffusing MPAct probe may lag behind, creating the appearance of a gradient. While an estimate of protrusion dynamics and diffusion rates argues against this scenario, we directly tested this potential alternative mechanism by treating migrating cells with a Jasplakinolide, Latrunculin B, and Y-27632 cocktail (JLY, (*30*)) to arrest actin dynamics, and then monitored whether the gradient equalizes. After stopping all actin dynamics, F-tractin positive protrusions remained stably depleted of MPAct (Fig. 2I) despite a preserved ability of MPAct to undergo lateral diffusion in the inhibited cells (Fig. S3G). Furthermore, expression of MPAct caused no observable change in migration rates, indicating the reporter is relatively inert (Fig. S3H).

Another interpretation of the observed MPAct gradient might be that there is a specific distance between the membrane and actin cortex in the front that is different in the back. However, we found that increasing the length or flexibility of the of the MPAct sensor by inserting a ~6 nm long helical linker (EAAAR)_8_ (*31*), or a longer 121 a.a. flexible linker (Fig. S3H, (*20*)), did not change the front-to-back gradient (Fig. S3H-N). Increasing the linker length did reduce its diffusion coefficients (Fig S3I), confirming that the reporter is detecting more F-actin for the same conditions. This argues that local differences in MPAct reporter signal reflect differences in the overall F-actin concentration in a zone proximal to the plasma membrane rather than a specific difference in the distance of an F-actin layer in different regions of a cell. Thus, we conclude that the MPAct signal gradient reflects an underlying back-to-front gradient of the density of membrane proximal F-actin density that is low in the front of migrating cells.

### Gradients with low MPAct in the front in different cells types and for different migration conditions

We next tested if MPAct gradients are a general feature of migrating cells beyond linear migration on fibronectin stripes. Markedly, individual HUVEC cells migrating within a monolayer showed similar gradients (Fig. 3A). Furthermore, both HeLa and RPE-1 migrating on uniform collagen substrates in 2D showed robust MPAct gradients (Fig. 3B-C). In addition, TIRF imaging of RPE-1 confirmed that there are strong MPA density gradients at the adhesion surface (Fig. 3C, Movie S3). Finally, using RPE-1 migrating in soft 3D collagen matrices (0.5 mg/mL), we observed a similar gradient of MPAct (Fig. 3D). We conclude that gradients in MPA density with low levels in the front are a ubiquitous structural feature of cells that migrate alone or collectively and also in cells that migrate on linear tracks, on 2D surfaces or in 3D matrices.

**Fig. 3.**
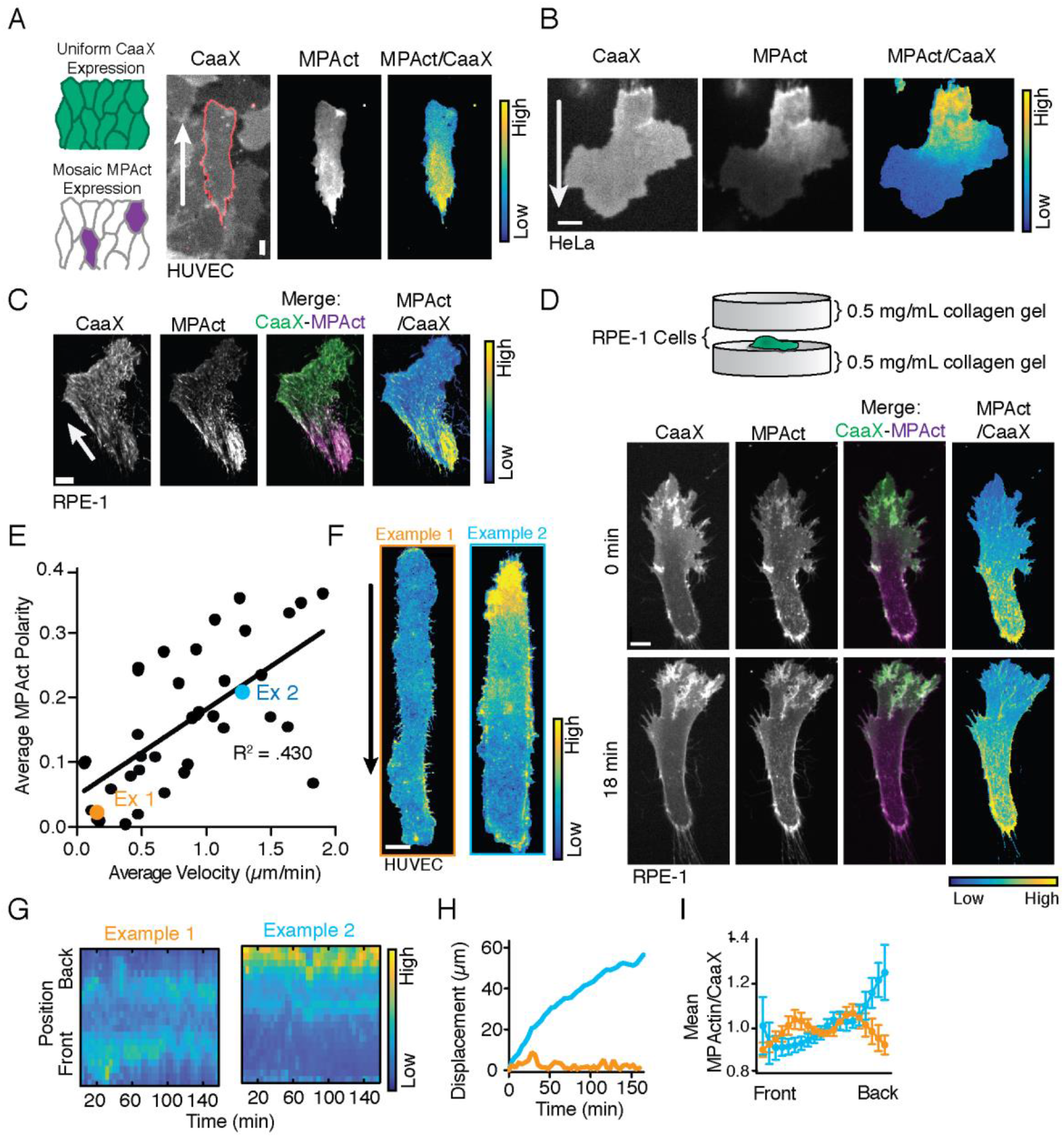
MPA density gradients are a general characteristic that distinguishes motile from non-motile cells. (**A**) Normalized MPAct/CaaX ratio in collectively migrating HUVEC monolayers with mosaic reporter expression. (**B**) Randomly migrating HeLa on collagen coated glass transiently transfected with MPAct-mRuby and iRFP-CaaX. (**C**) TIRF images of RPE-1 stably expressing MPAct-mRuby (magenta) and a CaaX-anchored sensor (green, as membrane marker) randomly migrating on uniform collagen coated glass. (**D**) RPE-1 cells stably expressing iRFP-CaaX (green) and MPAct-mCitrine (magenta) were plated between 0.5 mg/mL collagen matrix layers and imaged after 24 hours. (**E**) Mean MPAct polarity vs mean velocity for HUVEC migrating on 20 μm stripes for at least 1 hour. Mean MPAct polarity was calculated by subtracting the averaged back and front 10% of normalized MPAct signal. Velocity was calculated by taking the total displacement over time tracked. Line of best fit calculated using Prism 8 (GraphPad, San Diego, CA). n= 43 cells from 2 independent experiments. (**F, G**) Representative cells from 3E shown as two images (F) and kymographs (Left, orange and right, light blue). Analysis identical to 2B. Cells were tracked for > 140 min and imaged every 2 minutes. (**H, I**) For same cells as in 3F, centroid displacement over time (H) and quantification of average normalized MPAct/CaaX distribution (I). Error bars are standard deviation. For all images direction of cell migration is indicated by arrow. Scale bar is 10 μm.

We then returned to our initial hypothesis that low MPA in the front may weaken membrane retention in the front and thereby promote membrane protrusion and cell migration. Such a retention role of MPA suggests that the presence and steepness of the MPAct gradient directly correlates with a cell’s migration velocity. Indeed, when we plotted the relative MPAct polarity signal versus the average velocity we found a positive correlation between the two (R^2^ = .430, n= 44 cells from 2 independent experiments, Fig. 3E). Example cells show that cells with very low speed (Example 1) show a near symmetric distribution of MPAct/CaaX over time (Fig. 3F-I, orange) while migrating cells (Example 2) show a marked front-back asymmetry (Fig. 3F-I, light blue). Thus, gradients of MPA with low levels in the front is a general structural feature of migrating cells and polarity in MPA density correlate with cell velocity. We next set out to determine whether the gradient is merely the result of polarization and migration or whether it is induced concurrent with cell polarization.

### MPAct is locally decreased prior to protrusion in *de novo* polarization

Randomly migrating cells stochastically lose their polarization, before reorienting themselves and migrating in a new direction. Figure 4A shows an example of a change in direction of the MPAct gradient in a repolarizing RPE-1: Not only does the gradient reorient with the cell as it changes direction, but nascent protrusions in the new direction of migration (Fig. 4B, arrowhead, Movie S4) are already depleted of MPAct reporter signal. The dynamics of the directional change can be seen in individual cells using a kymograph analysis (Fig. 4C) and in a graphical representation of MPAct polarity and migration direction as a function of time (Fig. 4D). We conclude that MPAct gradients are established approximately in parallel with the development of a new global polarity and new direction of cell migration.

**Fig. 4.**
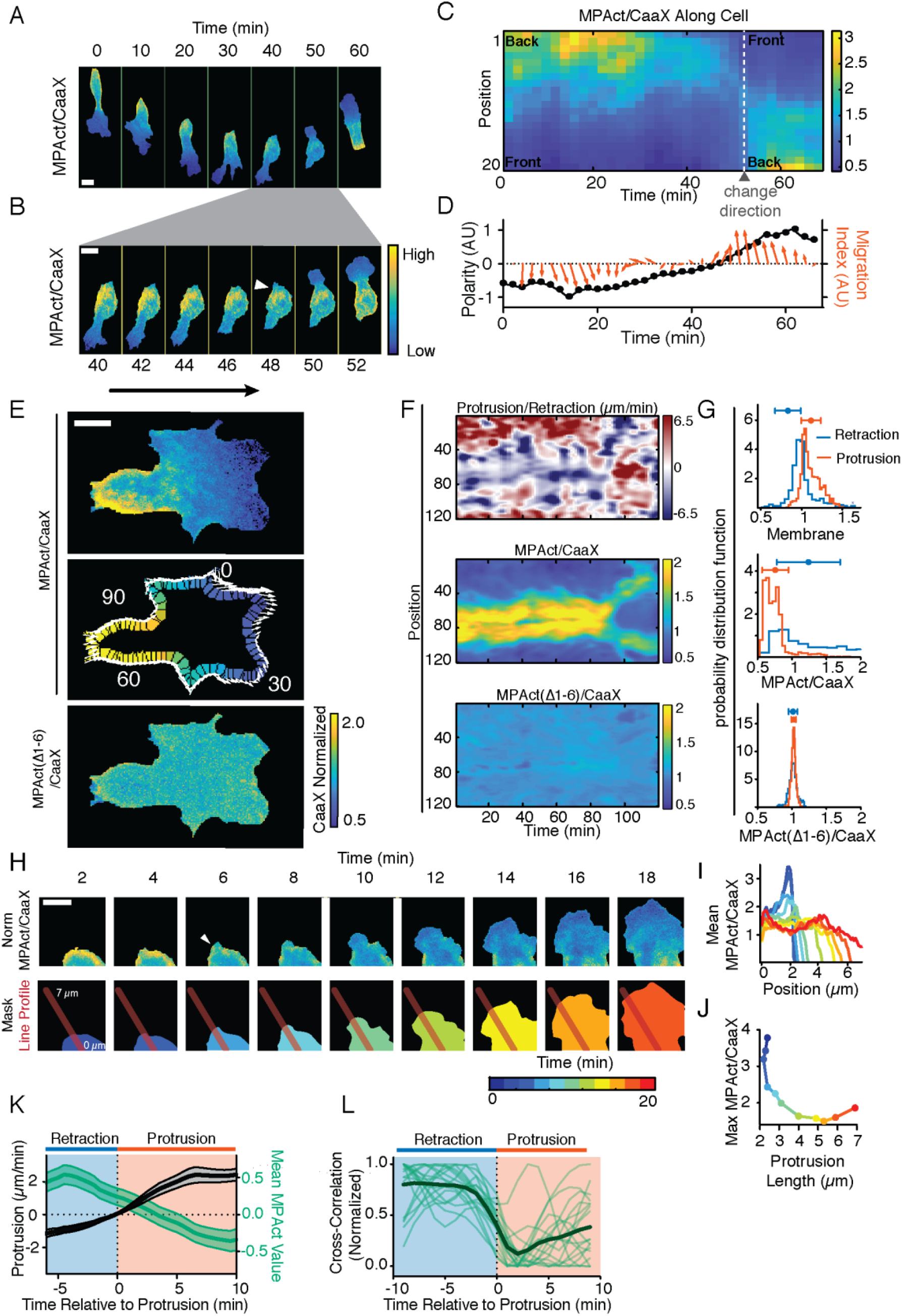
Global MPA density gradients are established concurrently with cell polarity. (**A-D**) RPE-1 co-expressing MPAct-mCitrine (magenta) and CFP-CaaX (green) migrating on uniform collagen (A) with repolarization event shown with higher time resolution (B), Kymograph analysis (C), and time course analysis of MPAct signal (black, left y-axis) and Migration Index (red, right y-axis) (D). Polarity represents average of peripheral positions 1-2 minus 19-20. Migration index shown as arrow marking the distance and direction traveled between subsequent time points (relative to future migration direction). (**E**) Control, comparing CaaX normalized MPAct-mCitrine to MPAct(Δ1-6)-mRuby polarity. Local protrusion direction and distance shown along the periphery by white arrows, global direction by black arrow. (**F**) Kymograph analysis for edge velocity (top), Normalized MPAct (middle), Normalized MPAct (Δ1-6) (bottom) versus time. Images taken every 1 minute. (**G**) Probability density function of normalized CaaX, MPAct and control MPAct (Δ1-6) comparing protrusive (red, top 20% of positive edge velocity) and retractive (blue, bottom 20% of negative edge velocity) windows. Mean and standard deviation of each distribution is indicated above. (**H-K**) (Top images) MPAct signal changes during local retraction and protrusion. Color codes used for analysis shown in masks below. Red line shows profile direction. (bar = 5 μm). (H). Line intensities for normalized MPAct profiles (I) and maximal MPAct signal changes in a nascent protrusion (J). Mean protrusion velocity and local MPAct signal change (K). Traces were aligned to the time of protrusion. Mean and 95% CI, n=25 repolarization events. **(L)**Alternative cross-correlation analysis in the front of HUVEC migrating on 20 μm stripes, n= 19 cells. Scale bar is 10 μm unless marked.

One prediction of our hypothesis that MPA inhibits membrane protrusions is that there should be a close local spatiotemporal anti-correlation between MPAct signals and local membrane protrusion. To test for such a local correlation, we parameterized the edge of randomly protruding cells into 120 evenly spaced regions, and subsequently measured the average normalized MPAct and local protrusion/retraction rates along the cell periphery (Fig. 4E, adapted from (*32*)). This analysis showed a close spatiotemporal anti-correlation between MPAct signals and protrusion (Fig. 4F, Fig. S4A-B). The control MPAct(Δ1-6) construct that does not bind F-actin showed no correlation, arguing that the correlation is indeed mediated by local MPA density differences (Fig. 4F, bottom, Movie S5). Interestingly, while the increase in local protrusion occurs in low MPAct areas (Fig. 4G and S4C, red), some retracting areas have low MPAct signals as well (Fig. 4G, and S4C, blue), suggesting that lowering of MPAct functions as a permissive instead of a causative mechanism for local protrusion.

Because membrane protrusions coincide with areas with low MPAct signals, we next determined whether membrane protrusions precede or follow a lowering of MPAct signals. By leveraging the frequent stochastic repolarizations of migrating RPE-1 (Fig. S4D-E), we found that prior to *de novo* protrusions there is a small but significant decrease in MPAct (Fig. 4H, arrowhead). This is particularly apparent in a line-scan analysis of the protruding region over time which shows MPAct signals start decreasing before the membrane extends (Fig. 4I-J). To more precisely analyze this delay, we used a window correlation analysis for cells that form a new local protrusion (example of a repolarization event in, Fig. S4D-E). After the specific region was selected, the edge velocity (Fig. S4F, bottom – black) and mean normalized MPAct (Fig. S4F, bottom – green) for that region was averaged and aligned *in silico* based on protrusion initiation (average speed > 0 μm/min). Across many repolarization events, this analysis showed that there is an approximately 4-minute delay between MPAct starting to decrease and the membrane starting to protrude (Fig. 4K). A similar result was also obtained using a cross-correlation analysis in stably migrating HUVEC, showing that normalized MPAct signal decreased between 3-4 minutes before protrusion (Fig. 4L). These lines of evidence suggest that local lowering of MPA density plays a key role in initiating membrane protrusions. Thus, this suggests that for cells to protrude, local MPA concentration must be sufficiently low to create a permissive state where cells can direct actin polymerization outward, protrude membranes and eventually generate a polarized cell front for migration.

### Rapid Rac-mediated local membrane protrusion is restricted to areas with low MPAct signals

Rac activity is required to trigger broad membrane protrusions at the front of many migrating cell types (*32*, *33*). We therefore tested whether Rac signaling and MPAct signals are spatiotemporally anti-correlated and if reduction in MPAct was able to confine Rac-mediated protrusion to a defined area. We used a Raichu-Rac1 FRET reporter (*20*) to assess the temporal and spatial relationship between Rac activity, MPAct signals and protrusion. Markedly, regions of high Rac activity and protrusion-retraction cycles were confined to areas of low MPAct (Fig 5A-B, Fig S5A-C). In a more precise analysis, we aligned the MPAct signal and membrane protrusion *in-silico* to the time of half-maximal Rac1 activation. Interestingly, this analysis showed that a MPAct signal reduction typically precedes the increase in Rac activity with MPAct signals then further decreasing along with the Rac activity raising to its maximal level. Given the close correlation, the initial drop in MPA density likely facilitates Rac activation and increasing Rac activity then further promotes a reduction in MPA density (Fig. 5C). Markedly, low MPAct signals are not sufficient to sustain Rac activity: Rac signals frequently decrease following a protrusion event while low MPAct signals remain refractory and persist for several minutes after the drop in Rac activity and onset of retraction (see arrows in Fig. 5A for periods where local Rac activity has dropped and an increase in the local MPAct signal is delayed by several minutes). Such a refractory mechanism was confirmed by treatment with JLY, which sustains the MPAct gradient while abrogating Rac polarity (Fig. S5 D-E). This suggests that low MPA has a role to increase the persistence of cell polarization for several minutes beyond a loss in a Rac activity gradient.

**Fig. 5.**
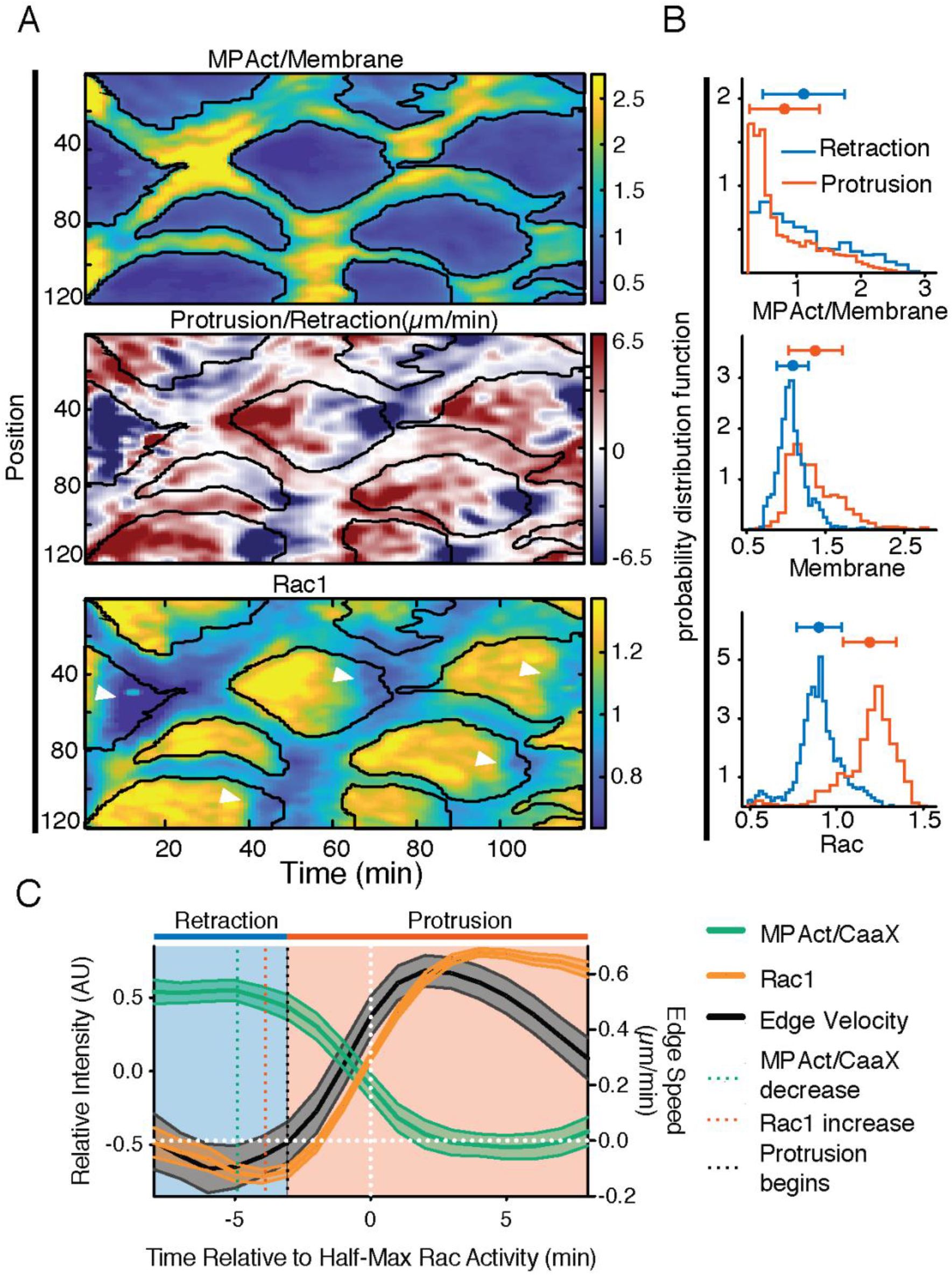
Rac1 and MPAct show high spatio-temporal anticorrelation. (**A**) Kymograph of normalized MPAct/CaaX (top), edge velocity (middle), and Raichu-Rac1 FRET (bottom) in RPE-1. Edge analysis identical to Fig. 4A. Black outlines marks areas of low MPAct signals. (**B**) Probability density function of MPAct (top), membrane (middle), and Rac (bottom) comparing top 20% (retraction, blue) and (protrusion, red) areas. Mean and standard deviation shown. (**C**) In silico activity alignment by half-maximal Rac activation, comparing protrusion (black, right y-axis), MPAct (green left y-axis), and Raicu-Rac1 (orange, left y-axis) signal time courses. Mean and 95% CI, n=30 events, 3 independent experiments.

To directly test the hypothesis that low levels of local MPA density are required for Rac-driven membrane protrusion, we used a Lyn-FRB and Tiam1-FKBP activation system to uniformly activate Rac (Fig. 6A (*34*)). In response to a rapamycin-triggered Rac activation, we tested whether different regions in the same cell with high or low MPAct have different protrusion responses. Indeed, rapid Rac activation triggered selective protrusions from particular peripheral sites before ultimately ending up with a broad radial protrusion (Fig. 6B-C). Consistent with a “biased protrusion” mechanism, the areas that had low MPAct signals were the first to protrude after rapamycin addition (Fig. BF, blue line), while the areas that had high MPAct remained initially unresponsive (Fig. 6B, red line). MPAct biased protrusion is evident upon close examination of Tiam1 stimulated cells (Fig. 6C-E). For the cell outlined in Figure 6C one can see in an example that the area initially devoid of MPAct (positions 1-20, Fig. 6D-E) rapidly protrudes soon after rapamycin addition, while the high MPAct areas fail to protrude for the first 5 minutes. We conclude that high levels of MPA are prohibitive for Rac-induced membrane protrusion and that lowering MPA levels is a required step that sensitizes sub-cellular regions for Rac-mediated F-actin polymerization and membrane protrusion.

**Fig. 6.**
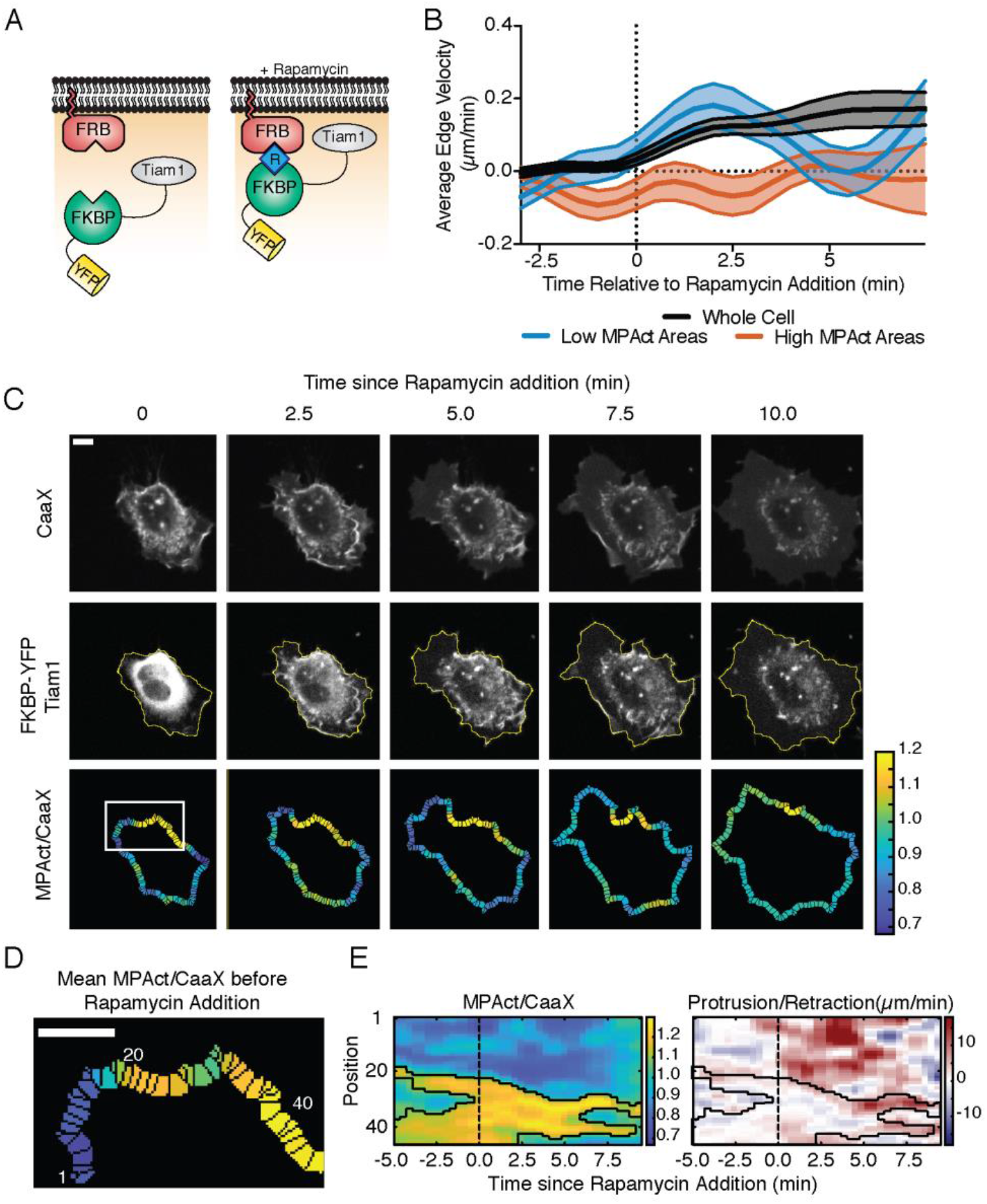
Low local MPA density biases nascent protrusion and promotes persistence. (**A**) Schematic of Tiam1-FKBP/Lyn-FRB Rac activation system. (**B**) Time course of edge velocity of the entire cell (black) comparing the 5% of the periphery with the highest (red) versus 5% of the lowest (blue) MPAct signal after uniform Rac activation. N= 27 cells, 2 independent experiments. (**C, D**) Example of cell response after uniform Rac activation. Membrane outline changes are shown as a yellow periphery overlaid on top of FKBP-YFP-Tiam1 image (C). Area indicated by box at the 0-time point is further analyzed in D-E. Scale bar is 10 μm. (D) shows parametrized area from (C) at time 0 with mean MPAct signal before rapamycin addition (**E**) Kymograph of normalized MPAct/CaaX (left) and edge velocity (right). Black outlines marks areas of low MPAct signals. Scale bar is 5 μm.

## Discussion

Our study reports the striking finding that MPA density has a strong back-to-front gradient with lowest levels in the front of both collectively migrating cells as well as in single cells migrating on linear stripes, on 2D surfaces or in a 3D environment. This was surprising since the cytosolic F-actin concentration is much higher in the front. We further showed that, under conditions where cells make one or more stochastic local membrane protrusions, such protrusions invariably occur from sites with locally low MPA density. Given that protrusions must push against membrane tension to move outward, our data suggests that local attachment between the cortex and membrane restricts local protrusions by creating high MPA density regions where membrane adhesion is strong, membrane tension is high, and protrusions are suppressed. Our data further suggests that local lowering MPA density can promote local Rac activation and local Rac activation further lowers MPA density to jointly promote local membrane protrusion. Our results with uniformly activated Rac further argues that biasing protrusion with low MPA density can serve as a cytoskeletal “memory” in the face of fluctuating migratory signals that maintains the low MPA density in the front for a few minutes and thereby enhances the persistence of cell migration. This provides a mechanistic interpretation of a previously described directional memory of moesin gradients observed in neutrophil-like cells (*17*).

A diverse range of molecular mechanisms are known to control F-actin near the plasma membrane including cofilin, which inhibits polymerization, severs actin filaments, and increasing the number of barbed F-actin ends (*35*). ERMs are primarily regulated by phosphorylation, phosphoinositides and adapters like EBP50 and E3KARP (*18*). Furthermore, there are additional actin-linked plasma membrane complexes and assemblies such as adherens junctions, integrins, clathrin and ER-plasma membrane junctions and capping proteins (*36*). The signaling mechanisms leading to the reduction in MPA density prior to protrusion may not be identical between cell types, as RhoGTPases and their effectors, capping and severing proteins, as well as Arp2/3 and formins, are known to have context-dependent roles in migration (*37*). Beyond the control of cell motility, the interface between membrane and cortex plays vital roles in exocytosis (*39*), generation of ER-PM junctions (*40*), diffusion-reaction dynamics kinetics at the membrane (*5*, *41*), and mechanotransduction (*42*). MPA density changes likely control specific steps in these processes and future studies are needed to explore the role of local MPA density in controlling other membrane localized processes regulated by cortical actin.

In conclusion, our study argues that the complexity of the signaling mechanisms controlling membrane protrusion and cell migration can be better understood by a framework where MPA density represents a fundamental structural parameter that integrates different signaling processes, directs local membrane protrusions and stabilizes cell polarity during cell migration. The MPAct reporter introduced here provides a powerful new tool for experiments aimed at understanding how different cells control membrane protrusions, directed cell migration and other membrane-localized and actin-regulated processes.

## Acknowledgments

We are grateful to the Meyer lab for helpful comments and discussions. We thank Yilin Fan and Nalin Ratnayeke for help with developing code and analysis techniques. We thank Olga Davydenko for help with the Leica TIRF system.

## Author contributions

AB and TM conceptualized the study and methodology, acquired funding, and wrote the manuscript. AB performed the investigation, validation, visualization, and data curation, and formal analysis of the data. AB and AH developed software for the analysis. AB, DG, AH and DC developed resources for the study. DG, AB, AH, and TM reviewed and edited the manuscript. TM supervised the study and administrated the project.

## Data and materials availability

All data, code, and materials used in this study will be made available upon request.

