## Supplemental Materials for "Membrane proximal F-actin restricts local membrane protrusions and directs cell migration"

**This PDF file includes:**

Materials and Methods
Table S1
Figs. S1 to S5
Captions for Movies S1 to S5

**Other Supplementary Materials for this manuscript include the following:**

Movies S1 to S5
Supplementary Reference (*43*)

### Materials and Methods

Cell culture.

HeLa cells (ATCC CCL-2), HEK293T, hTERT-immortalized HUVECs (HT-HUVECs as described in (*24*) hTERT-immortalized RPE-1 (ATCC CRL-4000), and rat mast cells RBL-2H3 were used to create all cell lines. HeLa, HEK293T, and RBL-2H3 cells were maintained in DMEM (Thermo Fisher, 11995-065) supplemented with 10% FBS (Sigma-Aldrich, TMS-013-B). HUVECs were maintained in EBM2 (LONZA CC-3156) supplemented with EGM2 (LONZA CC-4176). RPE-1s were maintained in DMEM-F12 (Life Technologies, 11039-047) supplemented with 10% FBS. Cells stably expressing fluorescent reporter constructs were created from RPE-1 or HUVECs via lentiviral transduction followed antibiotics selection and/or FACs sorting. Cells were passaged every 3-5 days to maintain sub-confluency

Antibodies and reagents

Phalloidin conjugated to Alexa Fluor568 (A12380), Hoechst 33342 (H3570) were from Thermo Fisher Scientific. CK-666 (SML0006), Bovine fibronectin (F1141), FITC-conjugated bovine collagen (C4361) were from Sigma-Aldrich. Y-27632 (13624) and Latrunculin A (ab428026) were from Cell Signaling Technologies. SMIFH2 (4401) from Tocris, bFGF (223-FB) from R&D Systems, Latrunculin B (144291) from Abcam, Jasplakinolide (sc202191A) from Santa Cruz Biotechnologies, PLL-PEG (PLL(20)-g[3.5]-PEG(5)) from SuSos and Bovine collagen (PureCol - 5005-100ML) from Advanced BioMatrix were also used.

DNA constructs and lentivirus production

Selected plasmid constructs and corresponding sequence information will be available on Addgene soon after publication. (http://www.addgene.org/Tobias_Meyer). All constructs generated here used Gibson assembly unless otherwise specified. Lentiviral backbones containing antibiotic resistance to neomycin, puromycin or blasticidin were created using pLenti-EF1a-MCS-IRES-antibiotic backbone. C1-YFP-CaaX and CFP-CaaX was previously described in (*19*). C1-Ftractin-mCitrine (FT-mCitrine) , C1-f-tractin-mTurqoise and the stoichiometric myosin II/F-actin activity reporter (pLV-Ftractin-muby3-p2a-MYL-mTurqoise-IRES-blast) was previously described in (*24*). GFP-moesin was a gift from the Bretscher Lab and have been described previously (*43*).

C1-MPAct-mCitrine was generated by PCR amplification of f-tractin *(23)* with a SDPPVAT linker to mCitrine to the CaaX Box of Kras4a (SGKKKKKKSKTKCVIM) with a SDPPVAT linker. C1-MPAct-mRuby was generated similarly. Lentiviral constructs were generated by digesting pLV plasmids with BamHI/EcoRI and inserting the full MPAct construct by PCR.

C1-FT'-mCitrine was created by digesting FT-mCitrine with HindIII and BamHI to remove the FT and replace it with a fragment of FT (residues 1-26, as described by (*23*)). C1-FT'-MPAct-mRuby was generated by digesting FT-mCitrine with NdeI/NotI-HF and using fragments created from PCRing part of the CMV promoter and the FT' fragment and mRuby-CaaX from MPAct-mRuby.

pLV-MPAct-mCitrine, pLV-MPAct-mRuby3 were generated by PCR amplification of C1-MPAct-pLenti-EF1a-MCS-IRES-neo or pLenti-EF1a-MCS-IRES-puro.

MPAct-$(EAAAR)_8$ – mCitrine was generated by ordering a codon optimized g-block and using gibson assembly with an EcoRI digested MPAct-mCitrine.

Lentivirus was generated in HEK293T cells plated in 10cm or 6 well collagen coated dishes. Prior to transfection media was switched from DMEM+10% FBS to Opti-mem. Cells were co-transfected with the third-generation packaging plasmids pMDLg/pRRE, pRSV-rev, pCMV-VSVG (provided by X. Liu, University of Colorado, USA) using Lipofectamine 2000, polybrene and a transfer vector containing the gene of interest. Viral supernatants were collected at 48-72 h post transfection then filtered using .22 µm vacuum filters (Fisher Scientisic, land concentrated using centrifugal filter units (100 kDa cutoff, Millipore, UFC910024).

Fibronectin Micropatterning for Linear Tracks

Fibronectin stamping was adapted from (*29*). Briefly, 20 µm linear tracks were generated creating a PDMS (Sylgard Silicone Elastomer Kit 184, ellsworth #4019862) stamp from an SU8 pattern. PDMS was cured for at least 3 hours at 65° C and then cut into smaller stamps (~0.5 $cm^2$) using a scalpel. 40 µL of 50 µg/mL fibronectin was spread onto the stamp and incubated at 37° C for at least 30 min. Stamps were washed 3x with 50 µl $ddH_2O$ and then left to air dry for ~15 minutes until stamps were completely dry. Stamps were then placed onto plasma cleaned Ibidi µ-Dish 35 mm, high dishes (Ibidi, 81156 , plasma treated for 30 seconds at 250 mTorr) using forceps. Stamps were then removed and incubated with 500 µL ,1 mg/mL PLL-PEG in 10 mM HEPES in $ddH_2O$ for 30 min at 37° C. Plates were washed 3x with 750 µL PBS and stored in PBS until use, usually within 1-2 weeks.

3D collagen gel migration

Glass bottom 96-well plates were plasma cleaned for 30 seconds at 250 mTorr. All collagen handling steps were performed on ice or with ice cold reagents until it was added to the plate. Advanced Biomatrix PureCol was diluted to a concentration of 2.4 mg/mL into 10x PBS pH 7.5. pH was adjusted to be 7.5 before further dilution into cold 1X PBS for a final concentration of 0.5 mg/mL. 60 µL of the collagen solution was added to the bottom of the well and allowed to polymerize for 2 hours at 37 degrees. The gel was washed with PBS 5x, always keeping the gel covered with some aqueous solution. 50 µL of RPE-1 cells at a concentration of $2 \times 10^4$ cells/mL were added to each well and allowed to settle for 2-3 hours before removing the 40 µL of media and adding an additional 60 µL of collagen on top the cells. Wells were washed with growth media 6 times and cells were imaged after 24 hours. Before imaging growth media was exchanged for extracellular buffer.

Live-cell imaging

Unless otherwise specified all migration assays occurred on glass-bottom (Cellvis, P96-1.5H-N), pretreated with 31 µg $ml^{-1}$ bovine collagen (Advanced BioMatrix, 5005-B) in PBS at 37 °C for at least 2 hours. 96 well plates were kept sealed during long term imaging. For 1D migration assays cells were plated on micropatterned ibiTreat 35 mm dishes as described above. Live cells were imaged in EGM2 buffered with 20 mM HEPES pH 7.4 or extracellular buffer (ECB, 125 mM NaCl, 5 mM KCl, 1.5 mM $MgCl_2$, 1.5 mM $CaCl_2$, 10 mM D-glucose supplemented with 1% FBS and 5 ng $ml^{-1}$ bFGF).

Images from Fig.3C were acquired on a Leica DMi8 S with Infinity TIRF equipped with 488 nm and 561 nm lasers and a 100x Plan Apo objective. 96 well plates were environmentally controlled using an Okolab stage top incubator (H301-K-Frame).

Images shown in Figs S1B, S2N, and analysis shown in S1A-D,S2C were captured using a fully automated fluorescence microscope (ImageXpress Micro XL, Molecular Devices), equipped with a Sola Light Engine (Lumencor), a Zyla 5.5 sCMOS camera (Andor), and using either a 10× objective and 0.3 NA (Fig S1, S2C) or a 20× 0.75 NA Plan Apo objective (Nikon, Fig S2L). When Hoechst was used for nuclear tracking it was incubated for 1 hour at a final concentration of 200 ng $ml^{-1}$ and imaged using a DAPI filter set.

All other images shown were acquired using a fully automated widefield/Yokogawa spinning-disc confocal fluorescence microscope system (Intelligent Imaging Innovations, 3i), with a Nikon Ti-E stand, equipped with Nikon 40× 1.3NA oil, 60× 1.27 NA water-immersion and 100x 1.4 NA oil objectives, an Olympus 60× 1.35 NA oil immersion objective, a 3i laser stack (405, 442, 488, 514, 561, 640 nm), a 3i ‘Vector’ photomanipulation device, an epifluorescence light source, (Sutter Lambda XL), a Yokogawa CSU-W1 scanning head with dual camera port, two sCMOS cameras (Andor Zyla 4.2), enclosed by an environmental chamber (Haison), and controlled by SlideBook 6.0 software (3i).

Transient transfections of cDNA or siRNA

For transient transfection of cDNA, 1-5 × $10^3$ cells per well were plated the day before transfection in glass-bottom 96-well plates coated with collagen as described above. On the day of transfection, 0.2 µg DNA of each construct (unless noted otherwise) and 0.25 µl Lipofectamine2000 (Life Technologies), diluted in 20-40 µL OptiMEM, was added following the manufacturer’s protocol. For HUVEC cells, the OptiMEM mixture was added to growth media up to a total volume of 100 µL. For all other cell types growth media was replaced with OptiMEM to a final volume of 100 µL. This transfection mix was replaced after 2-4 h with growth media and cells were imaged 16–24 h later. Pooled siRNA “Cherry-Pick” libraries were purchased from Dharmacon. For transfection, 1 x$10^4$ HUVEC per well were suspended in 80µL EGM2, and subsequently reverse- transfected with siRNAs to a final 20nM concentration with 0.25µL of Lipofectamine RNAiMAX (Thermo Fisher Scientific) in 20 uL of OptiMEM. The transfection mix was replaced with EGM2 after 8hrs. At 48hrs post transfection cells were analyzed.

JLY experiments

JLY freezing experiments were performed according to the protocol described in (*30*). Briefly, cells were pre-imaged in ECB, then incubated in 20 µM Y27632 for 10 min before the addition of 8 µM jasplakinolide and 5 µM latrunculin B.

1D FRAP acquisition and analysis.

For lateral diffusion calculations HeLa (Fig. 1I) or RPE-1s (Fig. S2D, S3I) were transiently transfected with plasmids as described above. Before imaging growth media was exchanged for ECB (described above). Confocal images of midsection of the cells were taken using a 60x oil objective. A 2 µm circle of lateral membrane was photobleached using consistent laser power. Identical bleaching parameters were used across experiments taken on the same day and the bleached area remained in the same location for each acquisition. For mCitrine or YFP family fusion proteins a 515 nm laser, CFP family proteins a 445 nm laser, and for RFP family proteins a 561 nm laster was used. For figure 1F images were acquired every second, For figure 1G images were taken at different frequencies over time (5×1s, 5×0.5s, 5×1s, 10×3x, 2-3×10s) to allow for proper fitting of the initial lateral diffusion and better estimation of the later time points. Photobleaching always occurred after the 5$^{th}$ frame (5 seconds). For experiments using Latrunculin A, a 2x solution (2 µM) dissolved in imaging buffer was kept at 37ºC and then incubated for at least 15 minutes before imaging.

For each FRAP curve, the membrane area was selected in MATLAB by hand using IMROI and then automatically segmented to mask out any background areas. For Fig. 1F, the background subtracted, bleach corrected based on the first 5 frames, and max-value corrected value in the membrane region was calculated for each time point. The fits shown in Fig. 1F were performed in Graphpad Prism using a one phase association For 1G, once the membrane was segmented, the region was auto-rotated such that it's longer axis ran along the X-axis. Then the average value for each x-value was generated for each time point. Line-scans were bleach corrected based on bleaching rates average value of the first 5 frames. The recovery of this area was fit to a 3D surface in MATLAB using cfit and the equation

$$F(x,t) = 1 - B \times \frac{r_o}{\sqrt{4Dt + r_o^2}} \times e^{-\frac{(x-c)^2}{4Dt+r_o^2}}$$

Where x is the distance in µm, t is time in seconds. We can solve explicitly for the initial bleach radius $r_o$, c is the offset of the bleach spot from 0, and D is the diffusion coefficient in $\mu m^2/s$. Rates were fit using the first 5 time points after recovery (.5 s each for 2.5 seconds).

2D FRAP acquisition and analysis.

To compare recovery rates between the front and back of polarized cells HUVEC (Fig. S2E) or RPE-1(Fig. S3F) expressing MPAct-mCitrine were plated on either a fibronectin stripes as described above or collagen coated glass bottom plate. Cells were monitored to make sure they were actively migrating and polarized. HUVEC cells were imaged using 40x-oil epifluorescence and RPE-1 cells were imaged using a 515 nm confocal laser. Images were taken at different frequencies over time (5 x 1s, 5x 0.5s, 5x 1s, 10x 3x, 2-3x10s). A 5 µm circle was FRAPed using a 515 nm laser in the same location after the 5$^{th}$ frame. Each cell was photobleached in the front and back, alternating between front or back being bleached first.

For analysis a mask was generated using the average difference between all the frames before and after photobleaching acquisitions from the same day to account for any deviations of the laser alignment. The average value in the mask was generated after background subtraction, bleach correction, and initial value normalization based on the first 5 frames. The fits shown in Fig. 2E were performed in Graphpad Prism using a one phase association. The individual $T_{1/2}$ shown in Fig. 2F and Fig. S3E are derived from those individual fits.

Cell tracking and motility analysis.

Segmentation of cell nuclei or membrane and tracking were adapted from previously designed MATLAB scripts (*24*). Masks of individual cells were identified using a modified Otsu's method or the adapthresh function in MATLAB . Cells that were not fully enclosed in the frame were excluded. Nearest-neighbor heuristics between subsequent frames was used to track cells. Cell velocity was determined on the basis of centroid displacements between subsequent time points (30 s- 5 min) averaged over 3 time points. Cell direction was determined by fitting the trajectory of the cell's XY location to a first order polynomial and then extrapolating the direction of the cell to minimize the effects of random centroid movements.

Sensor activity profiles in 1D migrating cells.

For HUVEC, MPAct-mCitrine and CFP-CaaX (Fig. 2A-D, 3E-I) or MPAct-mRuby and Rac1-Raichu FRET sensor (Fig. S3A-C) were stably expressed. Cells were plated at 2-3 × $10^3$ cells per 6-well Ibidi dish 24 hours later or a 1× $10^3$ . Time-lapse imaging of polarized and unpolarized cells was acquired using a Nikon Ti-E stand widefield-fluorescence microscope (Intelligent Imaging Innovations, 3i), a 40× 1.3NA objective (0.325 µm pixel resolution), acquiring mCitrine/CFP as appropriate at 1-4 min intervals. Cells were tracked over at least 60 minutes masked on the basis of the CFP-CaaX. The normalized MPAct value was generated by background subtracting each channel and then taking the ratio of the

YFP/CFP channels. RPE-1 stably expressing MPAct-mCitrine and iRFP-CaaX were plated similarly, but imaged using 515/641nm confocal lasers with the same objective.

Cells masks were tracked as described above and at a 3 µm ring was created around the periphery of the cell for each time point. The ring was then divided into 20 equally distributed segments along the front-back axis of the cell for better comparisons between cells. Each value of the normalized biosensor intensity in the ring mask was binned into one of the 20 segments based on its distance to the cell front thus creating the kymograph for each time point. For all sensor activity kymographs time is shown on the x axis, and each of the 20 spatial bins into a single line with front/back annotated on the graph.

Polarity Score and Average Speed

The polarity score was determined from activity sensor profiles in migrating cells. For Fig.3E, average polarity was calculated by using the absolute value of (mean(front 10%) – mean(back(10%)). All cells were tracked for at least 60 minutes with a 1-2 minute imaging interval. Average speed was calculated using centroid displacement over the time tracked. For Fig. 4A a similar calculations was done except it used the difference between mean(positions 1-2) – mean(position 19-20) for the cell shown fin Fig.4C at each time point.

Migration Index

The migration index shown in Fig. 4D is calculated by taking the centroid displacement between subsequent time points and representing the velocity and direction of cell movement by the length of the arrow and angle of the arrow respectively. Graphs were made using the quiver function in MATLAB.

Edge velocity and biosensor spatial activity maps

Analysis was adapted from (*32*). Briefly, time-lapse sequences were acquired at 15s - 2 min intervals, using a 40× 1.3 NA objective or 60x 1.27 NA objective and confocal illumination for RPE-1 cells. The cell body were defined based on a membrane signal (either CaaX or the YFP of the Rac1-Raichu FRET biosensor). For each frame, 120 markers were equally spaced along the cell boundary and shifted to minimize the median distance from the markers in the previous frame. This method enabled continuous tracking of markers and uniform spatial sampling of the cell edge. These markers were also used to create 120 bins in a 3 µm ring inside the cell. The average value of either the normalized biosensor signal was calculated for each bin such that the edge velocity and the sensor activity or distribution could be compared. A negative window velocity represents a retraction toward the cell body whereas a positive value represents a protrusion. For all edge velocity kymographs time is shown on the x axis, and each of the 120 windows is arrayed into a single line where window 1 and window 120 are adjacent. The windows are numbered such that they are always the same orientation with respect to field of view (e.g. if a cell moves at a constant angle, the protrusive region will show a high edge velocity within the same windows over time).

Biosensor activity histograms by edge velocity

For Figs. S4C,5C,6B the activity histograms were generated by analyzing single cell edge-velocity and activity maps. For each cell the spatiotemporal regions (both window number and time index) that were the most protrusive and retracting were selected by taking the 20% highest positive and negative instantaneous velocities values respectively. The normalized ratio (for MPAct constructs or CC-Rac1) or intensity (for CaaX and membrane constructs) for those corresponding points were selected and put into separate histograms.

Line-scan of MPAct and Protrusion Length

For the graphs in Fig. 4I-J F the normalized MPAct/CaaX ratio was generated after background subtraction and bleach correction of the individual channels in Fiji. A line segment shown in Fig. 4H (bottom, red line) was used to measure the intensity of the ratio with the plot profile function. The maximum value of the each profile was used for 4J. A mask of the cell was generated using thresholding of the CaaX channel and the protrusion length was determined by the overlap of the line segment that the mask.

Protrusion and MPAct Alignment

For all experiments, RPE-1 cells were plated sparsely on collagen coated 96-well glass bottom plates and analyzed with edge-correlation analysis scripts. Cells that showed a distinct repolarization based on edge velocity profiles were manually selected. The spatial region associated with the repolarization was automatically selected by thresholding for the top 20% of protrusion values and using the window numbers that are contained in that region. Once this region was selected, the corresponding MPAct values in that same region & time were used for comparison. MPAct values are normalized to be between -0.5 and 0.5 to allow for direct comparison between cells with different shapes and expression levels. The time average edge velocity was taken for this region to find the moment where the average velocity first became greater than 0 µm/s. This became the zero-point for protrusion and all average traces were aligned to this value. To be included in the final analysis, repolarization events had to have at least 10 minutes of tracking before and after the repolarization event.

Cross-correlation Analysis in 1D migration

Edge velocity and biosensor spatial activity mapping was performed on HUVEC migrating on 1D fibronectin stripes. For cells that were tracked for at least 1 hour, the protrusive area of each cell was automatically selected based on the windows with the highest edge velocity using a findpks function in MATLAB. The edge velocity and normalized MPAct in those windows was used to calculate a cross correlation based on an offset in time using Pearson's correlation as done in (*32*).

Rac1,Edge Velocity, & MPAct Alignment

RPE-1 Cells co-expressing the Rac1 FRET sensor and MPAct-mRuby were imaging using simultaneous confocal imaging. Areas of abrupt increase in Rac1 signaling automatically detected by segmenting areas of high Rac1 activity (normalized value > 1.1). The corresponding average Rac1, edge velocity and MPAct values were taken in those areas in a window of 10 minutes before and after the beginning of the Rac1 patch. All three average traces values were time aligned to ½-maximal activation of Rac1.

Tiam1 Edge velocity after Global Rac Activation

2-3 × $10^3$ HeLa were plated on collagen coated 96-well plates and then transfected 24 hours later with 100 ng of iRFP-CaaX and MPAct-mRuby, 270 ng Lyn-FRB, and 30 ng YFP-FKBP-Tiam1 (both described in (*34*)) sing .25 µL of lipofectamine in 20 µL of optimem. Media was changed after 2 hours and cells were imaged 24 hours later. Prior to imaging, growth media was exchanged for ECB. We found translocation was more robust when there was higher levels of Lyn-FRB expressed than FKBP construct. Translocation was induced by adding 100 µL of a 2x Rapamycin solution (500 nM) diluted into ECB. Cells were monitored using 40x confocal, imaging every 30 seconds. These cells were tracked and edge velocity analysis run as described above. The regions of highest and lowest MPAct was calculated based on a 5% moving average (6 windows) of the MPAct values prior to rapamycin addition. The average edge velocity for these windows were compared to that of the entire cell (all 120 windows)

**Supplementary Methods**
qPCR
For Figures S1A-B, RNA was isolated using the RNeasy Mini Kit (Qiagen, #74106) and then reverse transcribed using RevertAid First Strand cDNA Synthesis Kit (Thermofisher, K1621). Equal amount of cDNA were used in conjunction with iTaq Universal SYBR Green Supermix (Bio-Rad, 1725120) and amplified on a Roche LightCycler 480 Instrument II. Abundance was calculated relative to EEF1$\alpha$ after determining primer efficiency with a standard curve.

Phalloidin Staining
HUVECs (WT and MPAct-mCitrine expressing) cells were sparsely ($2.5 \times 10^3$/well) plated in separate wells in a collagen coated 96-well plate and allowed to adhere overnight. Cells were fixed with 4% paraformaldehyde. After 15 minutes the solution was washed 3x in PBS followed by 30 min incubation with a combined permeabilization/ blocking solution (0.1% Triton X-100, 10% FBS, 1% BSA, 0.01% $NaN_3$, in PBS). Phalloidin was diluted into permeabilization/blocking solution and incubated for 1 hour. Cells were washed 3x in PBS and imaged in PBS.

Comparison of speeds of WT and MPAct expressing cells, or after siRNA treatment
Movies were captured using a fully automated fluorescence microscope (ImageXpress Micro XL, Molecular Devices), equipped with a Sola Light Engine (Lumencor), a Zyla 5.5 sCMOS camera (Andor) using either a 20× 0.45 NA objective (Nikon) with DAPI and YFP filters at a 6 minute interval for 6 hours. Cells were tracked using the pipeline described in (*24*)**.**

For Fig. S1C-D cells were imaged 48 hours after siRNA treatment. Prior to imaging, cells were incubated at 200 ng $ml^{-1}$ for 1 hour to stain nuclei for tracking. For Fig. S2C WT HT-immortalized HUVEC ($2.5 \times 10^4$) were plated on collagen coated 96-well glass bottom plate and allowed to adhere for 24 hours. Cells were plated in a mosaic (1 MPAct cell: 10 WT cells). Prior to imaging, cells were incubated at 200 ng $ml^{-1}$ for 1 hour to stain nuclei for tracking. Cell traces were divided into WT and MPAct expressing cells based on YFP expression. Only cells that were traced throughout the video were considered for velocity analysis. Average velocity was calculated by taking the average speed between each set of successive time points.

Table S1. List of qPCR Primer Pairs

| Gene | Forward Primer (5'-3') | Reverse Primer (5'-3') |
|---|---|---|
| Moesin | GCCCGACAGTCCTAGCTAAA | CAAACTCCAGCTCTGCATCC |
| Ezrin | GATAGTCGTGTTTTCGGGGA | CTCTGCATCCATGGTGGTAA |
| Radixin | TGGAACGTCTAAAACAAATTGAAG | TCGACGCTCCTTTTCAAGTC |
| EEF1$\alpha$ | GATGGCCAGTAGTGGTGGAC | TTTTTCGCAACGGGTTTG |

Figure S1

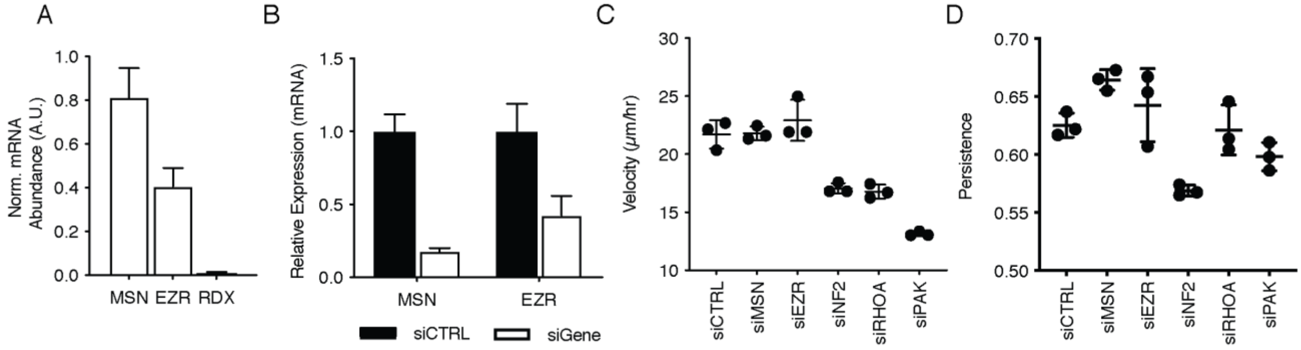


**Fig. S1 Knockdown of moesin or Ezrin has no significant effect on cell migration or persistence**

(**A**) Abundance of moesin, ezrin and radixin in WT hTert HUVEC by qPCR relative to EEF1a expression. (**B**) Knockdown efficiencies of pooled (20 nM final) siRNA against moesin and Ezrin in WT hTert HUVEC monolayers after 48 hours. (**C**) Mean velocity of individual HUVEC cells after 48 hours of siRNA treatment. Each point represents the mean velocity of a 96-well plate. Cells were imaged every 10 minutes for 16 hours and tracked using a script adapted from (*24*). (**D**) Mean persistence for individual HUVEC cells from (C). Each point represents mean persistence for all cells in the 96-well plate. Persistence is calculated by taking displacement/total distance traveled. For all experiments n = 3 technical replicates, Error bars are standard deviation.

Figure S2

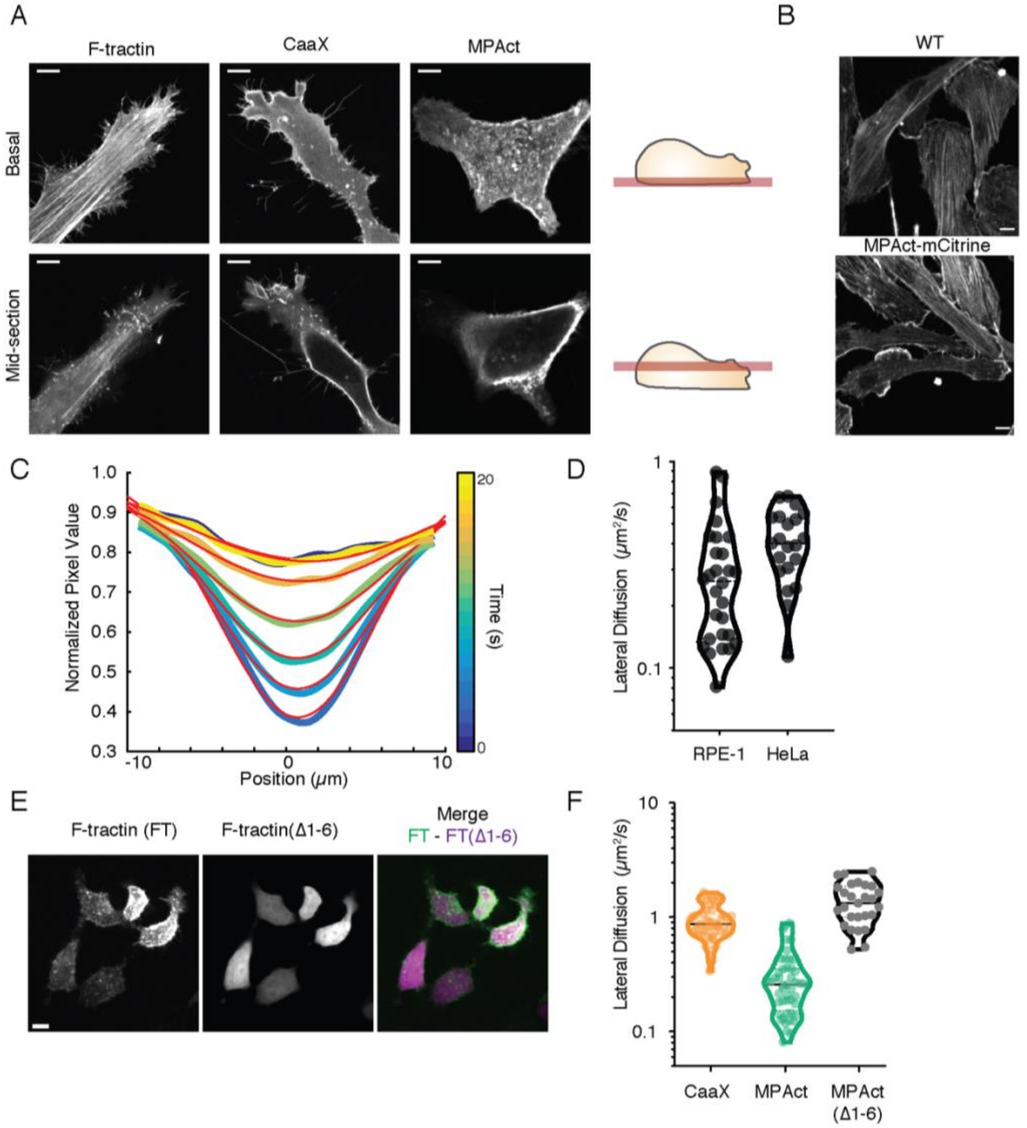


**Fig. S2** Control experiments characterizing the MPAct reporter

(**A**) Confocal slices of HeLa transiently transfected with F-tractin-mCitrine (left),YFP-CaaX (middle), and MPAct-mCitrine (right). Basal (top) and mid-plane (bottom) are shown. scale bar is 10 µm. (**B**) WT (top) and and MPAct-mCitrine (bottom) expressing hTert HUVEC fixed and strained with phalloidin-568. (**C**) Sample gaussian fitting of a single FRAP curve of CFP-CaaX. Position along the membrane (±10 µm) is shown in the X axis, time on the Y axis. Time is shown in a parula colormap with corresponding line of best fit overlaid in red. (**D**) Reproducibility in MPAct lateral diffusion between cell types. Lateral diffusion rates in the membrane were calculated in unpolarized HeLa (n = 20 cells) and RPE-1 (n=28 cells). (**E**) Max projection of RBL2H3 transiently co-expressing F-tractin-mCherry (left), F-tractin(Δ1-6)-mCitrine (middle). Dotted lines are quartiles. (**F**) Diffusion rates calculated from FRAP experiments preformed on the lateral membrane of RPE-1 based on 3D Gaussian profile fitting for CFP-CaaX, MPAct-mCitrine, and MPAct(Δ1-6)-mCitrine. n=51,48, and 27 respectively from 2 independent experiments. CFP-CaAX and MPAct-mCitrine are identical to Fig 1I.

Figure S3.

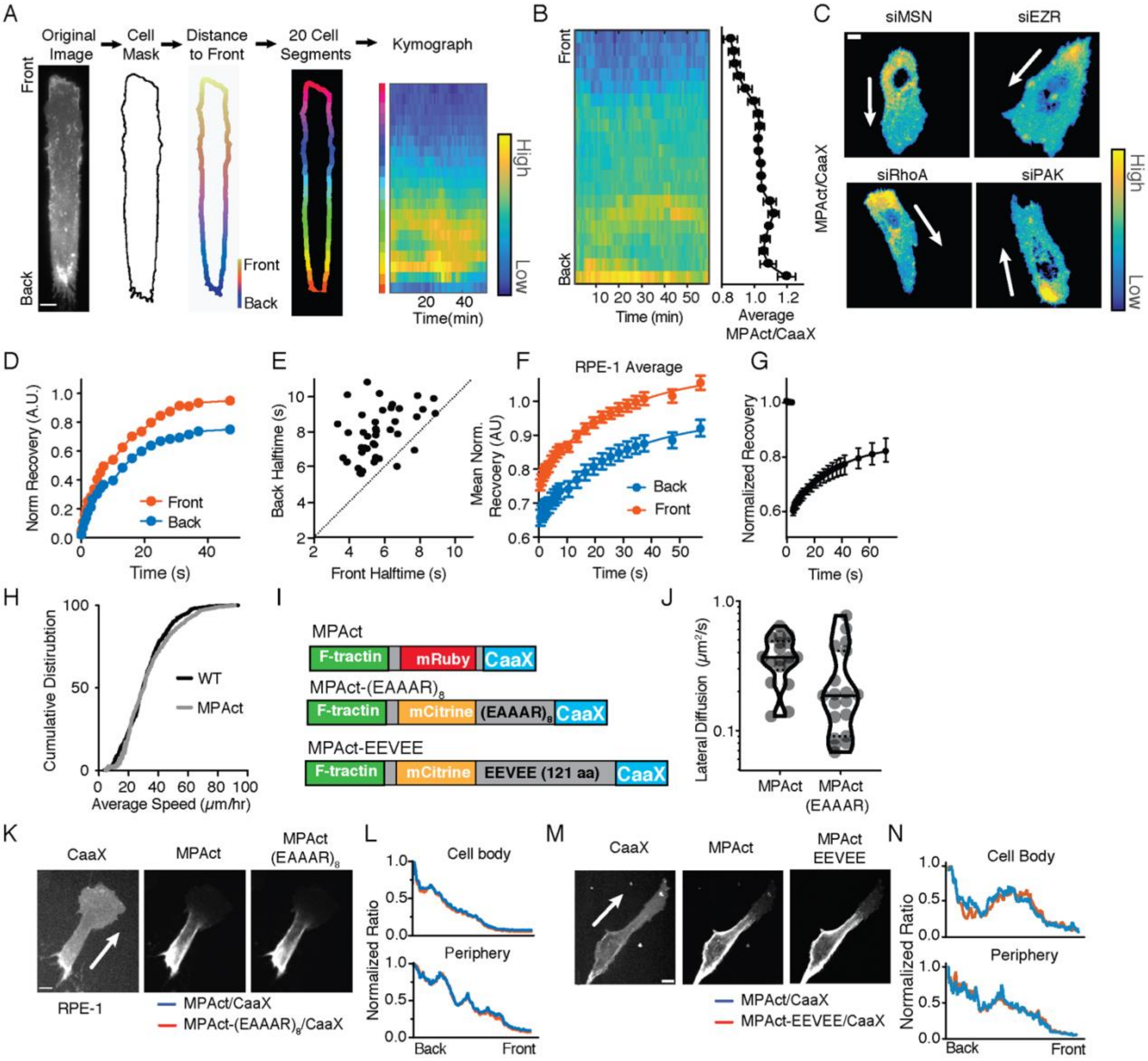

A
Original Image
Cell Mask
Distance to Front
20 Cell Segments
Kymograph
Front
Back
High
Low
20 40
Time(min)
B
Front
Back
10 20 30 40 50
Time (min)
0.8 1.0 1.2
Average MPAct/CaaX
C
siMSN
siEZR
siRhoA
siPAK
MPAct/CaaX
High
Low
D
Norm Recovery (A.U.)
Front
Back
Time (s)
E
Back Halftime (s)
Front Halftime (s)
F
RPE-1 Average
Mean Norm. Recvoery (AU)
Back
Front
Time (s)
G
Normalized Recovery
Time (s)
H
Cumulative Distirubtion
WT
MPAct
Average Speed (μm/hr)
I
MPAct
F-tractin
mRuby
CaaX
MPAct-(EAAAR)8
mCitrine
(EAAAR)8
MPAct-EEVEE
EEVEE (121 aa)
J
Lateral Diffusion (μm²/s)
MPAct
MPAct (EAAAR)
K
CaaX
MPAct
MPAct (EAAAR)8
RPE-1
MPAct/CaaX
MPAct-(EAAAR)8/CaaX
L
Cell body
Periphery
Normalized Ratio
Back
Front
M
CaaX
MPAct
MPAct EEVEE
MPAct/CaaX
MPAct-EEVEE/CaaX
N
Cell Body
Periphery
Normalized Ratio
Back
Front

**Fig. S3 Characterization of the MPAct gradient in migrating cells**

(**A**) Example gradient quantification during migration on fibronectin stripes. Cells are masked using a 3 µm ring around the cell perimeter. The MPAct signal in the periphery is normalized using the CaaX signal and segmented into 20 equal-length segments. The average value MPAct in each segment, is represented in the kymograph along the y-axis. (**B**) Kymograph of MPAct signal in RPE-1 stably expressing MPAct-mCitrine and iRFP-CaaX migrating on 20 µm FN stripes. Mean distribution over time shown on the right. (**C**) Ratio of MPAct-mCitrine/CFP-CaaX in mosaic HUVEC monolayers after 48 hours of siRNA transfection. Cells were additionally incubated with Hoechst for nuclear tracking which causes lower ratio signal in the nuclear area. (**D**) Example of an individual HUVEC FRAP recovery curves after photobleaching in the front and back. (**E**) Half-time recovery rates from individual cells plotted in Fig. 2F for direct comparison (n=48 cells from 3 independent experiments). (**F**) Average recovery curves after FRAP for MPAct-mCitrine in the front (red) and back (blue) of RPE-1 migrating randomly on uniform collagen. N=37 matched cells from 2 independent experiments. Line of best fit shown fits using 1 phase association and error bars are 95% CI. (**G**) Average recovery of FRAP of MPAct-mCitrine in the lateral membrane of RPE-1 after 20 minutes of JLY incubation. N = 22 cells from 1 experiment. (**H**) Cumulative distribution of the average single cell speed comparing randomly migrating cells in a mosaic of WT and MPAct-mCitrine expressing hTert HUVEC. WT N = 571 Cells, MPAct N = 290 Cells. (**I**) Schematic of linker length variants made for this study. MPAct-$(EAAAR)_8$ (middle) , inserts a 40 a.a. rigid helix in between the fluorescent protein and the CaaX. MPAct-EEVEE (below) , inserts a 121 a.a. flexible linker from the Raichu family of FRET biosensors in between the fluorescent protein and the CaaX. (**J**) Diffusion rates calculated from FRAP experiments preformed on the lateral membrane of RPE-1 based on 3D Gaussian profile fitting for MPAct-mCitrine and MPAct-$(EAAAR)_8$-mCitrine. n=18 &19 respectively from 1 experiment. Dotted lines are quartiles. (**K**) Confocal images of RPE-1 cells co-expressing iRFP-CaaX, MPAct-mRuby3, and MPAct-$(EAAAR)_8$-mCitrine. (**L**) Quantification of CaaX ratioed line-scans of MPAct-mRuby (blue) and MPAct-$(EAAAR)_8$-mCitrine(red) for the cell body(top) an periphery (bottom). (**M**) Confocal images of RPE-1 cells co-expressing iRFP-CaaX, MPAct-mRuby3, and MPAct-EEVEE -mCitrine. (**N**) Quantification of CaaX ratioed line-scans of MPAct-mRuby (blue) and MPAct-EEVEE-mCitrine(red) for the cell body(top) an periphery (bottom). Unless otherwise specified, cells in all panels are migrating on collagen coated 96-well plate. Images were background subtracted before ratio and measurements. For all images direction of cell migration is indicated by a arrows on the CaaX image. Scale bars, 10 µm.

Figure S4.

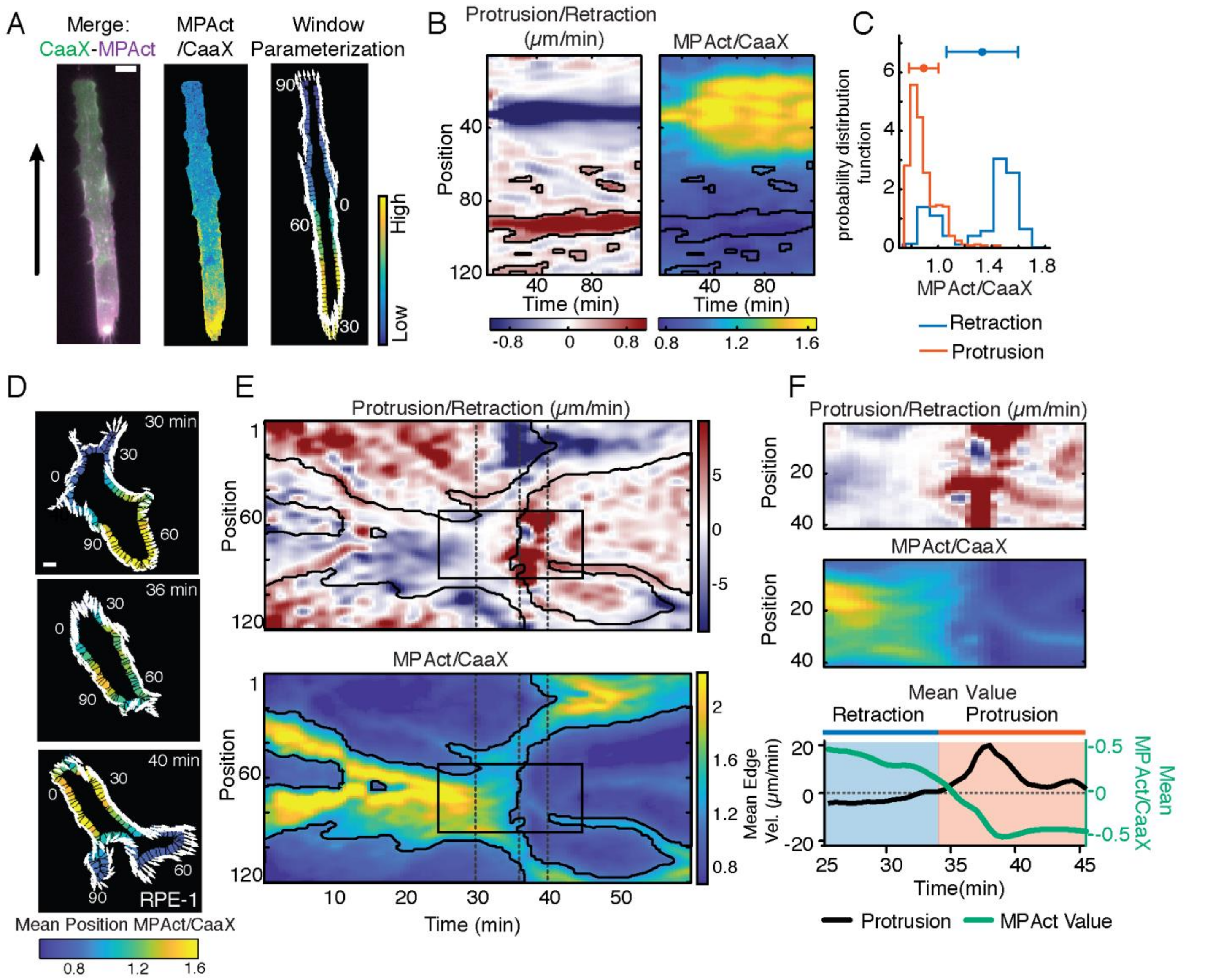
A
Merge: CaaX-MPAct
MPAct /CaaX
Window Parameterization
90
0
60
30
High
Low
B
Protrusion/Retraction (μm/min)
MPAct/CaaX
Position
40
80
120
40 80
Time (min)
-0.8 0 0.8
0.8 1.2 1.6
C
probability distribution function
6
4
2
0
1.0 1.4 1.8
MPAct/CaaX
Retraction
Protrusion
D
30 min
36 min
40 min
RPE-1
Mean Position MPAct/CaaX
0.8 1.2 1.6
E
Protrusion/Retraction (μm/min)
Position
1
60
120
5
0
-5
MPAct/CaaX
2
1.6
1.2
0.8
10 20 30 40 50
Time (min)
F
Protrusion/Retraction (μm/min)
Position
20
40
MPAct/CaaX
Mean Value
Retraction
Protrusion
Mean Edge Vel. (μm/min)
20
0
-20
Mean MPAct/CaaX
0.5
0
-0.5
25 30 35 40 45
Time(min)
Protrusion
MPAct Value

**Fig. S4. Alignment of local protrusion velocity and MPAct signal changes along the cell periphery**

(**A**) Edge velocity segment analysis for HUVEC expressing CFP-CaaX and MPAct-mCitrine migrating on 20 µm FN stripes. Analysis pipeline was identical to that used in Fig. 5A. Merge (left) and normalized MPAct (middle) are shown. For each segment, mean CaaX normalized MPAct (right) is indicated by the color of the region and edge velocity in the following frame is indicated by the direction and length of the white arrow emanating from that window. (**B**) Segment analysis for edge velocity (left) and Normalized MPAct (right) for the cell in A. Segment number (y-axis) vs Time (x-axis) psuedocolored to show protrusive areas (red) and retracting areas (blue) for protrusion and retraction, or with parula for the normalized MPAct. Black outline marks protrusive area (**C**) Distribution of normalized MPAct/CaaX values for segments that were in protrusive (red, top 20% of positive edge velocity) and retractive (blue, bottom 20% of negative edge velocity). Mean and standard deviation is indicated above. (**D**) Segment parameterization for the time points indicated by the dotted lines shown in Fig. S4E are shown to the right with segments labeled. For each segment, mean MPAct is indicated by the color of the region and displacement in the following frame is indicated by the direction and length of the white arrow emanating from that segment. (**E**) Edge correlation analysis for a single cell undergoing a re-polarization event showing edge velocity (above) and normalized MPAct (below). The 3 dashed vertical line show the time point shown in (D). Black box shows repolarization event that is examined more closely in Fig. S3F. Black outline shows area of low MPAct. (**F**) Kinetics of protrusion and MPAct changes during repolarization as outlined in Fig. S3E. Semi-automatic analysis of edge velocity maps selected the windows and timing of a single polarization. Edge velocity (top) and normalized MPAct/CaaX ratio (middle) for that region. Bottom shows average values for each vs time. Protrusion is shown in black (left y-axis) and MPAct is shown in green (right y-axis).

Figure S5

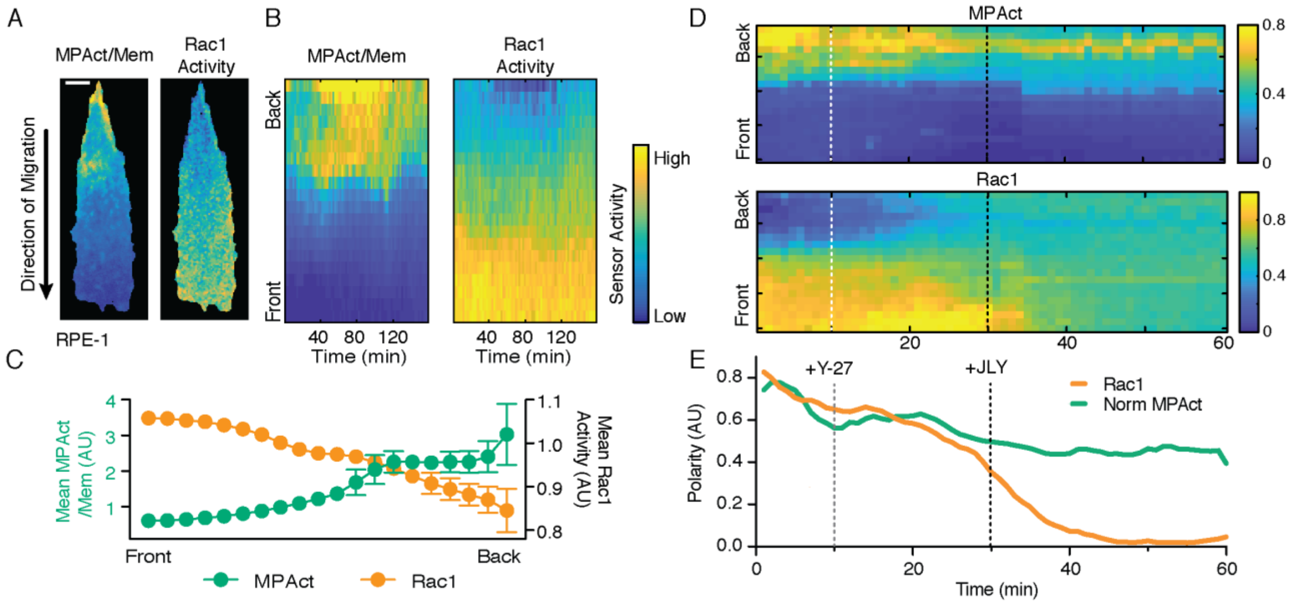


**Fig. S5 Rac1 activity is spatially anticorrelated with MPAct**

(**A**) RPE-1 stably co-expressing MPAct-mRuby (Left) and Raichu-Rac1 FRET sensor we plated onto 20 µm fibronectin stripes and imaged using confocal imaging with a CFP/YFP dichroic and two cameras enabling simultaneous acquisition of CFP/FRET signals. MPAct-mRuby and YFP(membrane control) were imaged with 561 and 515 nm lasers. Resulting normalized intensities are shown for a single time point. (**B-C**) Mean distribution over time for Rac1 activity (black, right y-axis) and Normalized MPAct/Membrane (green, left y-axis) for the cell shown in A-B over 160 minutes. (**D**) Spatial Rac1 and MPAct activity in a randomly migrating RPE-1 after addition of JLY cocktail. Cells were incubated with 20 µm Y27632 for 20 minutes and then the JLY was added for a final concentration of 20 µm Y27632, 8µm jasplakinolide, and 5µm latrunculin B was added. Cells were imaged every minute. (Confocal, 40x). (**E**) Average polarity of MPAct and Rac1 from the cell analyzed in (D). Polarity was calculated as in Fig. 4D for each time point.

**Movie S1.**
Migrating HUVEC monolayers expressing iRFP-CaaX (left,green) transiently transfected with GFP-moesin (middle, magenta) migrating on glass coated collagen. Scale bar 10 µm.

**Movie S2.**
Ratio of normalized MPAct-mCitrine/iRFP-CaaX in HUVEC migrating on 20 µm fibronectin stripes shown in parula color scale. Cells are from independent experiments. Scale bar 10 µm.

**Movie S3.**
TIRF of RPE-1 stably expressing CC-Rac1 FRET sensor (left,green) used as a membrane marker and MPAct-mRuby3 (middle, magenta). Cell is randomly migrating on uniform collagen coated glass plates. Scale bar 10 µm.

**Movie S4.**
Repolarizing RPE-1 transiently expressing iRFP-CaaX (left) used as a membrane marker and MPAct-mRuby3 (middle) and the normalized MPAct/CaaX ratio (right). Cell is randomly migrating on uniform collagen coated glass plates. Scale bar 10 µm.

**Movie S5.**
Normalized MPAct-mRuby3/iRFP-CaaX (left) and MPAct-(Δ1-6)-mCitrine/iRFP-CaaX (right) in the same cell. Cell is randomly migrating on uniform collagen coated glass plates. Scale bar 10 µm
